# Embracing native diversity to enhance maximum quantum efficiency of photosystem II in maize (*Zea mays* L.)

**DOI:** 10.1101/2024.08.12.604917

**Authors:** Sebastian Urzinger, Viktoriya Avramova, Monika Frey, Claude Urbany, Daniela Scheuermann, Thomas Presterl, Stefan Reuscher, Karin Ernst, Manfred Mayer, Caroline Marcon, Frank Hochholdinger, Sarah Brajkovic, Bernardo Ordas, Peter Westhoff, Milena Ouzunova, Chris-Carolin Schön

## Abstract

Sustainability of maize cultivation would benefit tremendously from early sowing but is hampered by low temperatures during early development in temperate climate. We show that allelic variation of subunit M of NADH-dehydrogenase-like (NDH) complex (*ndhm1)*, discovered in a European maize landrace affects several quantitative traits relevant during early development in cold climates through NDH-mediated cyclic electron transport (CET) around photosystem I, a process crucial for photosynthesis. Starting from a genome-wide association study (GWAS) for maximum potential quantum yield of photosystem II in dark-adapted leaves (Fv/Fm) we capitalized on large phenotypic effects of a hAT transposon insertion in *ndhm1* on quantitative traits early plant height (EPH), Fv/Fm, chlorophyll content and cold tolerance caused by reduced protein levels of NDHM and associated NDH components. Analysis of the native allelic series of *ndhm1* revealed a rare allele of *ndhm1* which is associated with small albeit significant effects on maximum potential quantum yield of photosystem II in dark- and light adapted leaves (Fv/Fm, ΦPSII) and early plant height compared to common alleles. Our work showcases the extraction of novel, favorable alleles from locally adapted landraces, offering an efficient strategy for broadening the genetic variation of elite germplasm by breeding or genome editing.

## Introduction

With more than 200 million hectares of maize harvested in 2022 (∼ 13 % of global arable land; FAO, 2022), enhancing the sustainability of its cultivation presents a significant opportunity to improve the environmental impact of agriculture worldwide. Earlier planting of maize increases resource efficiency with respect to available nutrients and water while simultaneously avoiding leakage of nitrogen into ground water. Further, the introduction of herbicides into the environment can be decreased by earlier soil coverage and fast early development through increased competition with weeds (Wezel et al., 2014; Nouri et al., 2022). Early sowing dates lead to earlier soil coverage which benefits resource efficiency of maize cultivation and increases biomass accumulation by extending the vegetative period and avoiding summer drought during flowering (Frei, 2000; Kucharik, 2006; Parker et al., 2017). However, the earlier sowing occurs, the higher is the risk for maize plants of encountering cold temperatures in the initial stages of development which can have adverse effects on various physiological parameters (Lainé et al., 2023). It has been shown that cold stress decreases biomass accumulation due to impaired photosynthetic parameters such as net CO_2_ assimilation, chlorophyll content and maximal quantum efficiency of photosystem II (Fv/Fm; Leipner et al., 1999; Burnett and Kromdijk, 2022; Lainé et al., 2023). Thus, genetic improvement of maize early development for an earlier soil coverage is a promising approach to increase sustainability of global agriculture.

Development of genetically improved crops builds on genetic variation for traits of interest originating either from standing variation within a crop and its close relatives, or from introduction of new variation through biotechnology (Liang et al., 2021). Introduction of new genetic variation can be untargeted by randomly generating mutations, or by directly targeting candidate genes (Lorenzo et al., 2023; Marone et al., 2023). Untargeted approaches do not require *a priori* knowledge about biological processes underlying trait expression, but the frequency of favorable mutations is generally low, thus requiring large screening populations to identify desirable phenotypes (Liu et al., 2020; Lorenzo et al., 2023). Targeted approaches on the other hand require information about the genes underlying the traits of interest, which is often not available for important quantitative traits. In addition, engineering of alleles that outperform their native counterparts requires in-depth knowledge about the functional mechanisms underlying the targeted process, especially in complex biological networks. Here, we demonstrate the identification of a biological process that influences traits of interest by a large-effect quantitative trait locus (QTL) followed by extensive analysis of allelic variation at the locus which revealed a beneficial allele for early development and photosynthetic traits.

In a doubled-haploid (DH) library derived from three landraces Mayer et al. (2020) mapped haplotype-trait associations for a number of quantitative traits related to early plant growth. In their phenotypic evaluation of landrace-derived DH lines in 11 different field environments, minimum temperatures ranged from -6°C to 3.5°C imposing considerable cold stress on the plants during early development (Hölker et al., 2019). They identified several QTL, associated with early plant height (stages V4 and V6, EPH), potentially linked to biomass accumulation under cold stress.

Building on the results of Mayer et al. (2020) we investigated the functional consequences of native allelic variation at a QTL on maize chromosome 2 with strong effects on EPH and the photosynthesis-related parameters potential maximum quantum efficiency of photosystem II (Fv/Fm) and chlorophyll content in the field. Our objectives were to functionally characterize the effects of the QTL on the three traits, analyze allelic diversity of candidate genes in the population and explore, if trait expression is temperature dependent. By genetic and functional characterization of the QTL for EPH, Fv/Fm and chlorophyll content we pinned down *ndhm1* as causal gene with pleiotropic effects on all three traits. We identified a rare structural variant of *ndhm1* in the landrace Kemater which is associated with improved EPH and Fv/Fm compared to more common variants and could show that a defective allele at the *ndhm1* locus leads to strong growth depression during early development which is enhanced by cold temperatures.

## Results

### Fine-mapping of an early development QTL

In a GWAS involving three European landraces, Mayer et al. (2020) identified QTL for EPH. We extended the GWAS with additional traits by measuring the maximum potential quantum yield of PSII (Fv/Fm) and chlorophyll content (SPAD) described in Hölker et al. (2019) using non-overlapping 10-SNP windows as haplotype markers. We observed a shared genomic region linked to EPH and photosynthetic traits Fv/Fm and SPAD (Table S1, Figure S1). This target region of 249 kb on chromosome 2 (Chr2:23,083,888-23,333,363; B73_AGPv4) was defined based on overlapping confidence intervals of four QTL for target traits EPH (V6), Fv/Fm (V4, V6) and SPAD (V3; Table S1). In three of the four QTL the same 10-SNP haplotype window showed the strongest association to target traits, thus we defined it as the lead haplotype (Chr2:23,328,380-23,333,363; B73_AGPv4, Figure S1B). The lead haplotype had an environmentally stable negative effect on EPH, Fv/Fm and chlorophyll content.

A flow-chart describing material development for fine-mapping is shown in Figure 1A. Two DH lines (KE0482, KE0678) from the landrace population in which the GWAS was performed were crossed to create a bi-parental population. They were selected to segregate at the QTL on chromosome 2 but to have similar genomic backgrounds based on genome-wide SNP markers (Figure S2). The two DH lines showed phenotypic segregation during early development, KE0678 exhibited significantly lower EPH and had lighter green leaves compared to KE0482 (Figure 1B, Figure S3), but did not differ significantly from each other for final plant height, lodging, tillering and flowering time (Figure S3). F_2:3_ recombinant inbred lines (RILs) were derived from the cross of KE0482 and KE0678 and phenotyped in a field experiment. In the F_2:3_ RILs, EPH and leaf color exhibited a significant association with SNP markers in the target region on chromosome 2 but not with the other nine maize chromosomes (Figure S4). Heterozygous F_2:3_ RILs showed an intermediate phenotype between the midparent value and RILs homozygous for the KE0482 allele indicating partial dominance (Figure S5).

**Figure 1:**
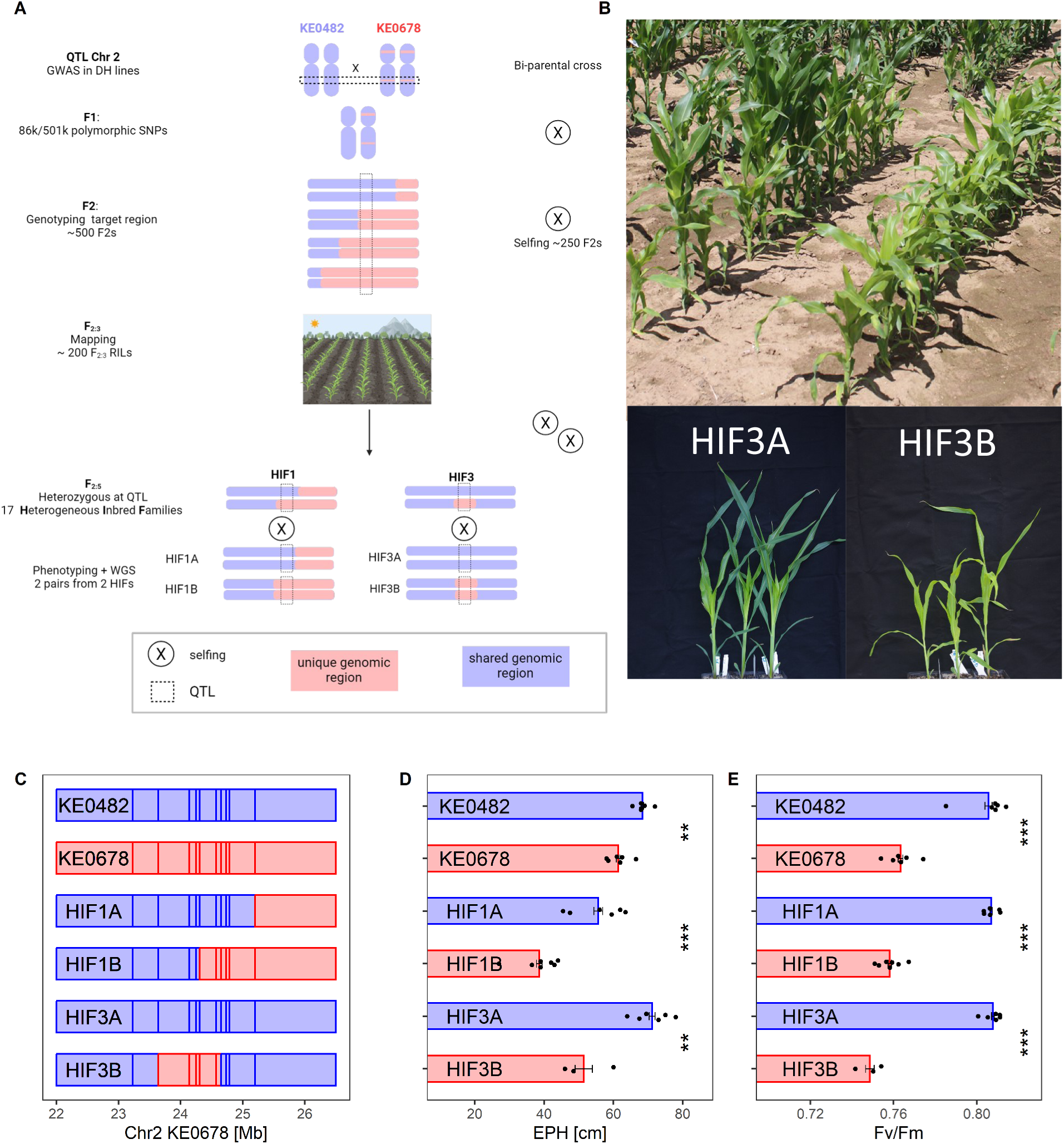
Fine-mapping of a QTL associated to early plant height (EPH) and Fv/Fm in growth stage V4. A) Scheme for fine-mapping of a QTL for EPH and Fv/Fm on maize chromosome 2. Genomic fragments unique to KE0678 are colored red B) Top: Phenotypic segregation during vegetative growth of F_2:3_ RILs linked to the QTL on chromosome 2. Bottom: HIF3A and HIF3B in a growth chamber experiment in optimal conditions C) Genetic composition of key heterogeneous inbred families (HIFs) that enabled fine-mapping to a 314 kb genomic region. Vertical lines represent genomic positions of KASP markers from Table S5. Genomic fragments inherited from KE0678 are colored red. D,E) Phenotyping of key HIFs for EPH (D) and Fv/Fm (E) in a growth chamber. Bars show means ± SE (n = 3-6 plants) and dots observations from single plants. Significant differences (t-test) are marked with stars. \*\*\**P* < 0.001, ** *P* < 0.01. Figure 1A was created with BioRender.com

Heterogeneous inbred families (HIFs) were developed by selfing selected F_2:3_ RILs from the KE0482xKE0678 cross for three more generations to construct pairs of lines differing for the QTL allele within a near-isogenic background. In growth chamber experiments, HIF1A and HIF3A resembled KE0482 phenotypically for traits EPH and Fv/Fm, while HIF1B and HIF3B resembled KE0678 (Figures 1B, 1D, 1E). For precisely defining the fine-mapped region overlapping between HIF1B and HIF3B, we assembled high contiguity genomes of KE0482 and KE0678 using PacBio long reads and re-sequenced the four HIFs (HIF1A, HIF1B, HIF3A, HIF3B) with Illumina short reads. Using SNPs derived from re-sequencing data mapped against the KE0678 genome assembly we defined the genomic region associated with EPH and Fv/Fm as a 314 Kb genomic fragment, covering six gene models (Figure 1C, Figure 2A, Figure 2B, Table S2).

**Figure 2:**
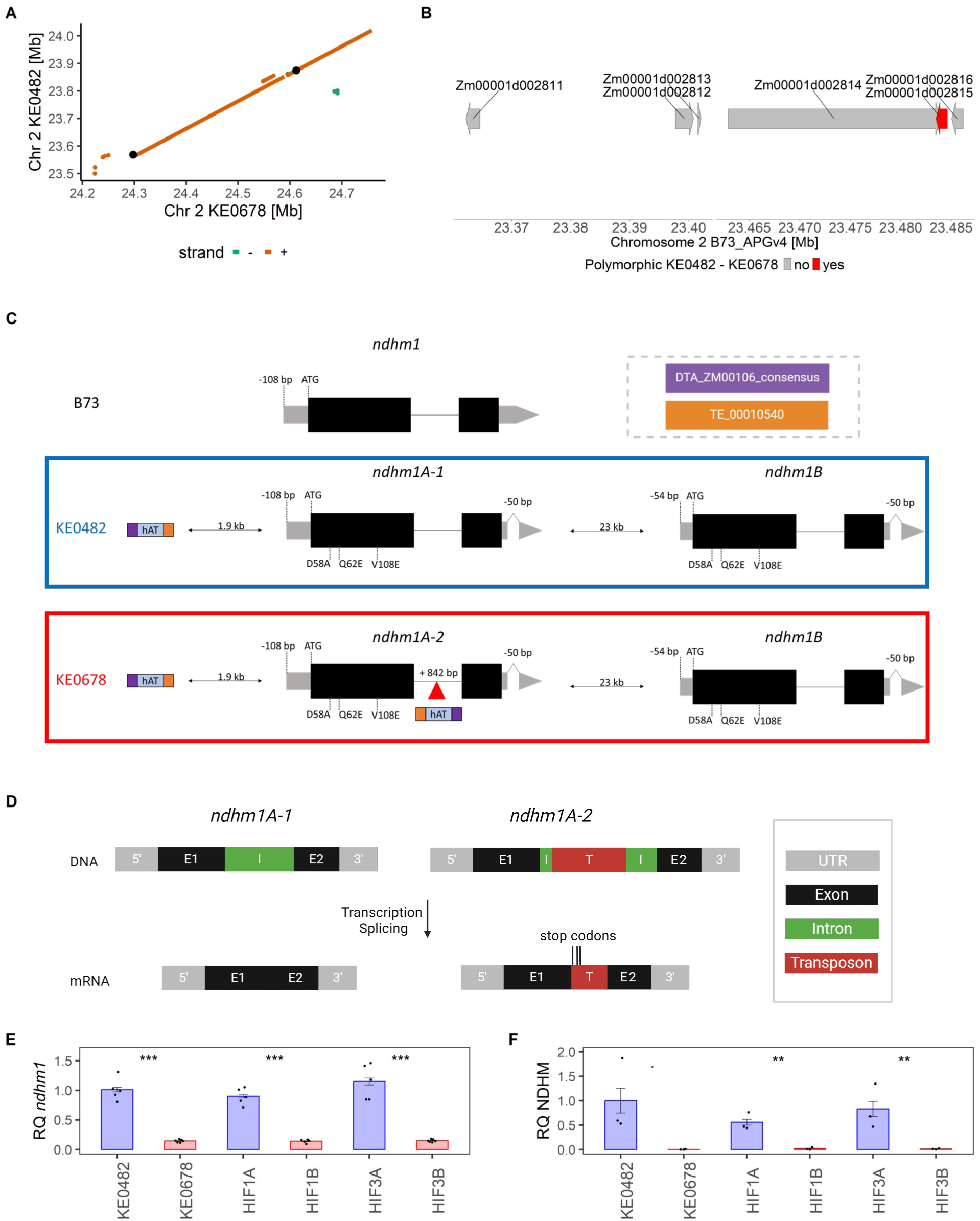
Candidate gene analysis in fine-mapped region. A) Pairwise sequence alignment (Sequence identity > 99 %, length > 1kb) of the fine-mapped region in KE0482 and KE0678. Black dots: Flanking markers of the fine-mapped region. B) Genes in fine-mapped region annotated in B73_AGPv4 reference genome. Red: *Zm00001d002815* (*ndhm1*) is polymorphic between KE0482 and KE0678. C) *Ndhm1* alleles and their polymorphisms in comparison to B73 reference sequence of KE0482 and KE0678. In *ndhm1A-2* a hAT transposon, flanked by two TIR motifs (purple, orange) is inserted in the intron (red triangle) D) Transcripts derived from *ndhm1A-1* and *ndhm1A-2*. E, F) Relative transcript (E) and protein (F) levels of *ndhm1* in KE0482, KE0678, HIF1A, HIF1B, HIF3A and HIF3B. Bars show means ± SE (n = 3 plants). Significant differences (t-test) are marked with stars. \*\*\**P* < 0.001, \*\**P* < 0.01. *P < 0.1.* Figure 2D was created with BioRender.com

The fine-mapped 314 kb genomic region is highly colinear between KE0482 and KE0678 (Figure 2A). Only one polymorphism overlapping with a gene model was observed between KE0482 and KE0678, a transposon insertion in the intron of a homologue of *Zm00001d002815,* making it the most likely candidate gene underlying the QTL (Figure 2B, 2C). *Zm00001d002815* codes for subunit M of the plastid NADH-dehydrogenase-like complex (NDH), central in NDH-mediated cyclic electron transport (CET) in plants (Shen et al., 2022). According to maizegdb.org two loci in B73 code for NDHM, *Zm00001d002815* (Chromosome 2) and *Zm00001d025952* (Chromosome 10). The amino acid sequence of the NDHM locus on chromosome 2 is highly conserved in the 26 US NAM assemblies with only 5/210 variable amino acids positions (Hufford et al., 2021). In contrast, the amino acid sequence of the chromosome 10 locus is weakly conserved and compared to *Zm00001d002815* sequence identity is 30% and 50% in the two alternative protein sequences for *Zm00001d025952* and includes large gaps (File S1, File S2). Thus, the locus on chromosome 10 likely does not function as NDHM and we call *Zm00001d002815* and its homologues on chromosome 2 in other maize genome assemblies *ndhm1*. Compared to B73, KE0482 carries a *ndhm1* copy with four amino acid exchanges and several SNPs and InDels in the 5’- and 3’ untranslated region (UTR) including a 50 bp deletion in its 3’ UTR (Figure 2C *ndhm1A-1*). All amino acid exchanges, as well as the UTR variations are common variants in the US NAM assemblies (Hufford et al., 2021), thus we assume this copy to be functional (File S3, S4). Additionally, we observed the insertion of a hAT transposon 1.9 kb upstream of *ndhm1A-1,* which is not present in B73 (Figure 2C). The hAT transposon insertion is found upstream of the respective *ndhm1* copy in KE0678 (*ndhm1A-2)* as well. An identical 842 bp hAT transposon insertion is found in the intron of *ndhm1A-2* of line KE0678 and results in the transcription of additional sequence originating from the hAT transposon causing three premature stop codons (Figure 2C-D, Figure S6A, File S3). Due to the premature stop-codons the resulting amino acid sequence lacks the second exon of NDHM, leaving it likely non-functional considering the high degree of sequence conservation of NDHM (File S3). A second copy of *ndhm1*, named *ndhm1B*, is present in both DH lines (Figure 2C) and is identical for KE0482 and KE0678. The *ndhm1B* sequence is identical to the *ndhm1A-1* 3’ UTR, exon and intron sequence up to a breakpoint 54 bp 5’ of the start codon. No sequence similarity to *ndhm1A-1* and *ndhm1A-2* is observed in the sequence located 5’ of this breakpoint. Thus, *ndhm1B* misses half of the 108 bp 5’ UTR region annotated in B73_AGPv4, impacting the putative promoter sequence and further 5’-located cis-regulatory elements. The sequence present in *ndhm1A-1* and *ndhm1A-2* but not in *ndhm1B* overlaps with the transcription start site in the shoot of maize reference line B73 and with a transcription factor (TF) binding site (File S5, File S6; Mejía-Guerra et al., 2015; Savadel et al., 2021).

### Functional validation of *ndhm1* as candidate gene for the early development QTL

*Ndhm1* expression in whole leaves was reduced to about 14% on transcript and 1% on protein level in KE0678, HIF1B and HIF3B compared to KE0482, HIF1A and HIF3A (Figure 2E, 2F, Figure S7). The remaining NDHM protein in KE0678, HIF1B and HIF3B is either translated from a transcript derived from *ndhm1A-2* where the hAT insertion is spliced from the mRNA or from *ndhm1B*. Transcripts specific for *ndhm1B* were amplified from KE0482 and KE0678 cDNA, indicating that at least some transcriptional activity remains (Figure S6B).

NDH is a protein complex consisting of at least 29 subunits, organized in five different subcomplexes which form a super complex with PSI for CET (Shen et al., 2022; Zhang et al., 2024). It has been shown in maize that null mutants of different NDH subunits affect accumulation of the whole NDH complex (Zhang et al., 2024). Leaf protein levels of all 16 NDH subunits and alternative CET pathways (PGR5/PGR1LA) were reduced in lines with the transposon insertion in n*dhm1A* (Figure 3A, Figure S8, Figure S9). An exception is plastid terminal oxidase 2 (PTOX2), where expression is almost doubled in lines with *ndhm1A-2* while levels of PTOX1 do not differ based on the *ndhm1* alleles. PTOX can remove electrons from the electron transport chain by oxidizing PQH_2_, thus upregulation of PTOX2 might indicate abnormalities in the photosynthetic electron transport caused by reduced accumulation of NDH components (Messant et al., 2024; Zhang et al., 2024). We conclude that the transposon insertion in *ndhm1A-2* reduces NDHM protein levels, leading to reduced accumulation of proteins required for NDH assembly.

**Figure 3:**
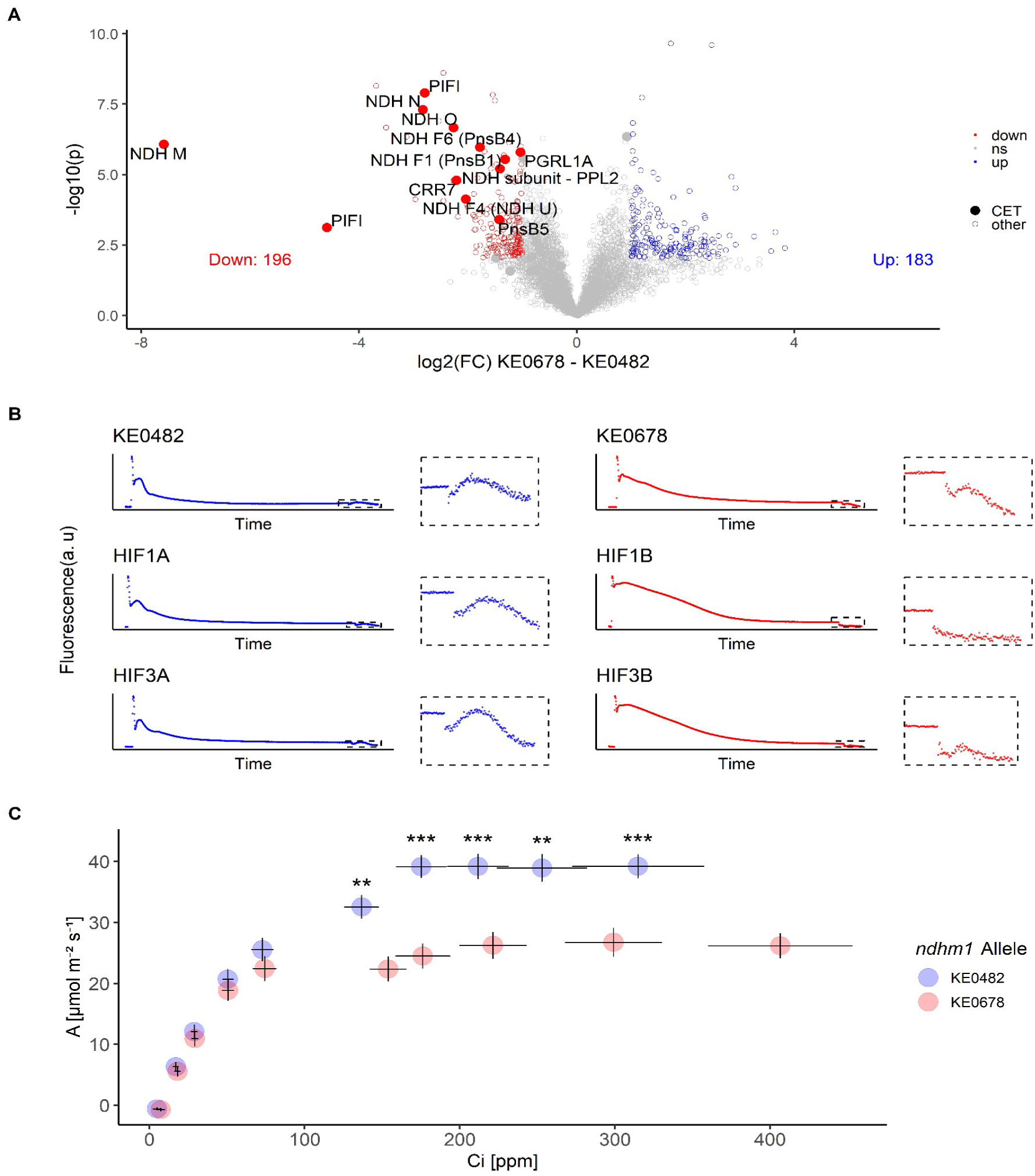
Impact of a hAT transposon insertion in *ndhm1 (ndhm1A-2)* on NDH components and photosynthetic parameters. A) Proteome data of 6 genotypes (KE0482, KE0678, HIF1A, HIF1B, HIF3A, HIF3B) in optimal conditions (N = 3 plants/genotype). Grouping was done by *ndhm1* allele (*ndhm1A-1 =* KE0482 allele or *ndhm1A-2* = KE0678 allele). One dot is one protein, filled dots are proteins taking part in cyclic electron transport. Cut-off for differential expression: log2(FC) > 1; FDR <5%. Proteins which are differentially expressed and involved in cyclic electron transport (CET) are labelled in the volcano plots. B) Chlorophyll fluorescence induction curves of dark-adapted plants. The small boxes and the right side of the plots show a zoom in the F0-rise indicative for CET after turning off a weak actinic light. C) Relationship between assimilation rate and intracellular CO_2_ concentration of KE0482, KE0678, HIF1A, HIF1B, HIF3A, HIF3B. Adjusted means were calculated for the two different *ndhm1* alleles considering the genomic background as random factor in a mixed linear model for each tested ambient CO_2_ concentration [Ca] separately. Dots are adjusted means ± SE. Significance was assessed by Wald tests for each Ca level independently. *** P<0.001, ** P<0.01

To investigate functionality of NDH-mediated CET *in vivo,* we recorded chlorophyll fluorescence traces of dark-adapted plants which can indicate NDH-mediated CET around PSI. The NDH-specific post-illumination fluorescence rise was strongly reduced in lines with the *ndhm1A-2* allele (Figure 3B). CET contributes significantly to ATP production required in CO_2_ fixating tissues of C_4_ plants, thus net carbon assimilation rates are dependent on a functional NDH complex (Ishikawa et al., 2016; Zhang et al., 2024). Net carbon assimilation rates at ambient CO_2_ concentrations of 400 ppm and higher were significantly reduced in genotypes with *ndhm1A-2* in the field (Figure 3C). Our results indicate that the hAT transposon insertion in *ndhm1A-2* in KE0678, HIF1B and HIF3B leads to an NDH-deficient phenotype significantly affecting target traits EPH and Fv/Fm.

We screened Mutator (Mu) transposon insertion lines from the BonnMu collection (Marcon et al., 2020) in the genomic background of inbred line F7 (Haberer et al., 2020; Win et al., 2024), carrying a single copy of *ndhm1* and identified an allele with Mu transposon insertion in the 5’ UTR of *ndhm1* (Figure 4A). The Mu transposon insertion is located 24 bp upstream of the ATG between the putative TSS and the start codon. The Mu transposon insertion leads to reduction of transcript levels by factor 10 compared to the wildtype (Figure 4B). Target traits EPH and Fv/Fm, were significantly reduced in the Mu induced mutant compared to its wildtype, similar to the effect of the hAT transposon insertion in *ndhm1A-2* in the line KE0678 (Figure 4C-D). The NDH-specific post-illumination fluorescence rise in the Mu induced mutant matches the observations from *ndhm1A-2* in the Kemater population indicating impaired NDH-dependent CET (Figure 4E, F).

**Figure 4:**
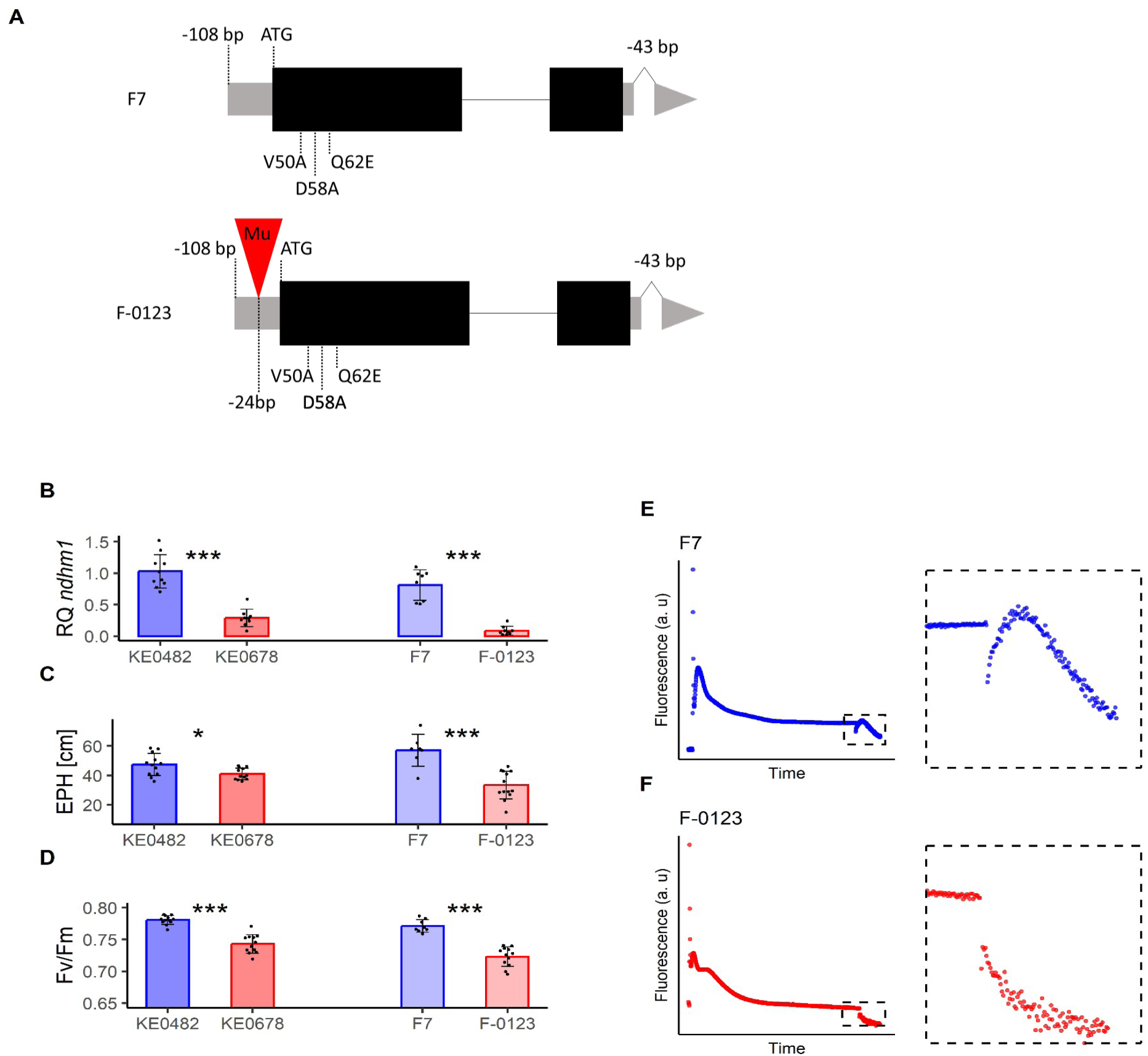
Phenotypic analysis of a *ndhm1* Mu-insertion mutant. A) Structure of *ndhm1* allele of Flint line F7 and derived Mu-insertion mutant F-0123 from the BonnMu collection compared to B73_AGPv4 reference sequence. The position of the Mu element (red triangle) 24 bp upstream of the start codon was determined by Sanger sequencing. B) Relative differences of *ndhm1* transcript levels for parents of bi-parental mapping population (KE0482, KE0678) and F7 and F-0123. C-D) Phenotypic differences for target traits EPH (C) and Fv/Fm (D) between F7 and F-0123 and the parents of the bi-parental mapping population. Bars show means ± SE. Significant differences (t-test) are marked with stars. *** P<0.001, ** P<0.01, * P<0.05. E, F) Chlorophyll induction curves for F7 and F-0123, respectively.

### Allelic diversity of *ndhm1* in the landrace Kemater

To assess if the landrace Kemater harbors alleles improving target traits we investigated the allelic series of *ndhm1* within the landrace. As a proxy for *ndhm1* alleles in Kemater we used the 10-SNP lead haplotype alleles from GWAS which is based on high-density genotypic data from 471 DH lines for which we had phenotypic data available (Mayer et al., 2022). The haplotype consists of ten consecutive SNPs from the 600k Axiom™ Maize Genotyping Array (Details on haplotype construction: Mayer et al., 2020) and does not overlap with *ndhm1* but is located 150 kb before *ndhm1* (lead haplotype: Chr2:23,328,380-23,333,363, *ndhm1:* Chr2:23,483,486-23,484,612; B73_AGPv4). This is because there are no polymorphic 600k SNPs between KE0482 and KE0678 in a 676 kb window starting 153 kb before and ending 523 kb after *ndhm1*. Four distinct haplotype alleles with allele frequencies between 6% to 44% were identified in the Kemater population and were used for grouping phenotypic data (Table S3). We extended our comparison of allelic effects with phenotypic data for traits EPH, Fv/Fm, chlorophyll content and final PH in 471 Kemater DH lines (Hölker et al., 2019). Strong phenotypic differences were observed between lines with the KE0482 haplotype and KE0678 haplotype for *ndhm1*, reflecting the results from the bi-parental population (Figure 5A-D). The KE0678 haplotype of *ndhm1* reduced EPH, Fv/Fm, chlorophyll content and final PH compared to the Hap1 and Hap2 groups as well, however to a smaller extent than compared to the KE0482 haplotype and not significant for EPH. In addition, a smaller but statistically significant increase in EPH, Fv/Fm, chlorophyll content and final PH was observed for lines with the KE0482 haplotype for *ndhm1* compared to lines with Hap1 and Hap2 (Figure 5A-D).

**Figure 5:**
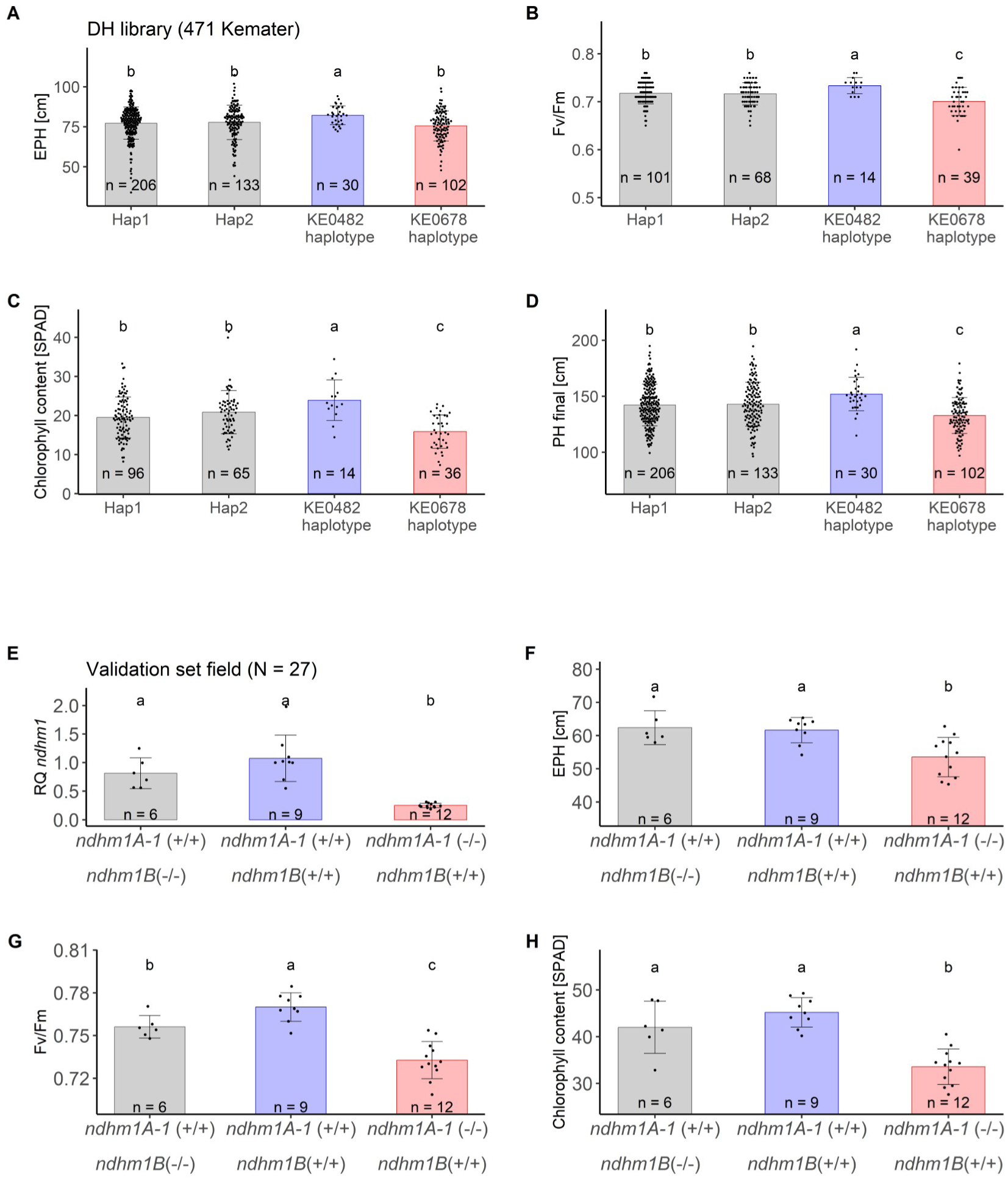
Analysis of allelic diversity of *ndhm1* in Kemater DH lines. A-D) Phenotypic data from 471 Kemater lines with high-density genotypic and phenotypic data grouped by their haplotype allele of the lead haplotype from GWAS. Blue: haplotype of KE0482 which carries a duplication of *ndhm1 (ndhm1B)* and no transposon insertion in *ndhm1 (ndhm1A-1).* Red: haplotype of KE0678 which carries a duplication of *ndhm1* (*ndhm1B*) and a transposon insertion in *ndhm1* (*ndhm1A-2*). Each dot represents the adjusted mean of one Kemater line in up to eleven combinations of environments and years for EPH (A), Fv/Fm (B), chlorophyll content (C) and final plant height (D). Significant differences of alleles are indicated by letters (lsd test, P < 0.05). E-H) Subset of 27 Kemater lines that was selected to be balanced for different *ndhm1* haplotypes for in-depth molecular characterization. Lines were genotyped for presence/absence of the transposon insertion in *ndhm1A-2* and presence/absence of *ndhm1B* and phenotyped for their differences in *ndhm1* transcript levels (E) and for target traits EPH (F), Fv/Fm (G) and chlorophyll content (H). Bars show means ± SE. Significant differences (lsd test) are indicated by letters.

For validation of the results on allelic diversity that were based on the lead haplotype from GWAS in proximity of *ndhm1* we selected 27 Kemater lines with a higher frequency of KE0482- and KE0678-associated *ndhm1* haplotypes as observed in the Kemater population. The selected Kemater lines were genotyped for presence or absence of the hAT insertion in *ndhm1A* and the presence or absence of *ndhm1B* (Figure S10, Figure S11, Table S4). 20/21 Kemater lines that had a second copy of *ndhm1* (*ndhm1B (+/+*)) carried the KE0482 or KE0678-associated *ndhm1* haplotype alleles. Only Kemater lines with the KE0678-associated haplotype carried the hAT transposon insertion in *ndhm1A (ndhm1A-1 (-/-))*. These results indicate that the KE0482- and KE0678 associated *ndhm1* haplotypes both are in LD with a duplication of *ndhm1*, while the hAT transposon insertion in *ndhm1A-2* is in LD with the KE0678 haplotype. *Ndhm1* transcript levels were strongly reduced in 12 Kemater DH lines with *ndhm1A-1*(-/-), supporting the notion that the insertion is causal for differences in expression levels (Figure 5E). *Ndhm1* transcript levels were slightly elevated in lines with the allelic combination *ndhm1A-1*(+/+)/*ndhm1B*(+/+) compared to lines with *ndhm1A-1(+/+)*/*ndhm1B(-/-),* however not significant (Figure 5E). On phenotypic level, EPH, Fv/Fm and chlorophyll content were significantly reduced by the hAT insertion in *ndhm1A* compared to all other lines (Figure 5F-H). The allelic combination *ndhm1A-1(+/+)/ndhm1B(+/+)* led to a significantly increased Fv/Fm compared to lines with *ndhm1A-1(+/+)/ndhm1B(-/-)* (Figure 5G). Results from the validation set show the same tendencies for investigated traits as observed in the full set of DH lines, however not all differences were significant.

To confirm that KE0482 and KE0678 *ndhm1* haplotypes are associated with a second copy of *ndhm1* (*ndhm1B*), we analyzed whole-genome sequencing data which was available for 15 Kemater DH lines from which three overlapped with the “Validation set field”. We assessed the copy number state of *ndhm1* by mapping short reads of the 15 lines against the B73_AGPv4 reference genome which has a single copy of *ndhm1*. In four DH lines, carrying either the KE0482 or KE0678-associated *ndhm1* haplotypes, normalized read depth exceeded two for most of two exons of *ndhm1*, indicating a duplication (Figure 6). A drop of the normalized read depth by half can be observed within the 5’ UTR of *ndhm1,* matching the breakpoint in the 5’ UTR of *ndhm1B* in KE0482 and KE0678 relative to B73. The remaining eleven DH lines with other *ndhm1* associated haplotypes display a normalized read depth of about one, without a drop of read depth in the 5’ UTR of *ndhm1*, indicating single-copy alleles of *ndhm1*. For four of the 15 Kemater lines we had PacBio high-contiguity genome assemblies available in addition to the short-read sequencing data (KE0095, KE0109, KE0482, KE0678). KE0095 and KE0109 both carry only one copy of *ndhm1*, validating results from the read depth analysis (File S3). The insertion of a hAT transposon 1.9 kb upstream of *ndhm1A-1* and *ndhm1A-2* in KE0482 and KE0678 (Figure 2C) was not present in KE0095 and KE0109. We expanded our investigation of allelic diversity of *ndhm1* to diverse maize lines by examining the presence of *ndhm1B* in the US maize NAM assemblies via BLAST (Hufford et al., 2021). We found a combination of *ndhm1A-1* and *ndhm1B* identical to the KE0482 variant in Tx303 but no allele similar to *ndhm1A-2* was detected (Zm00041ab076830 and Zm00041ab076840, File S3, S4).

**Figure 6:**
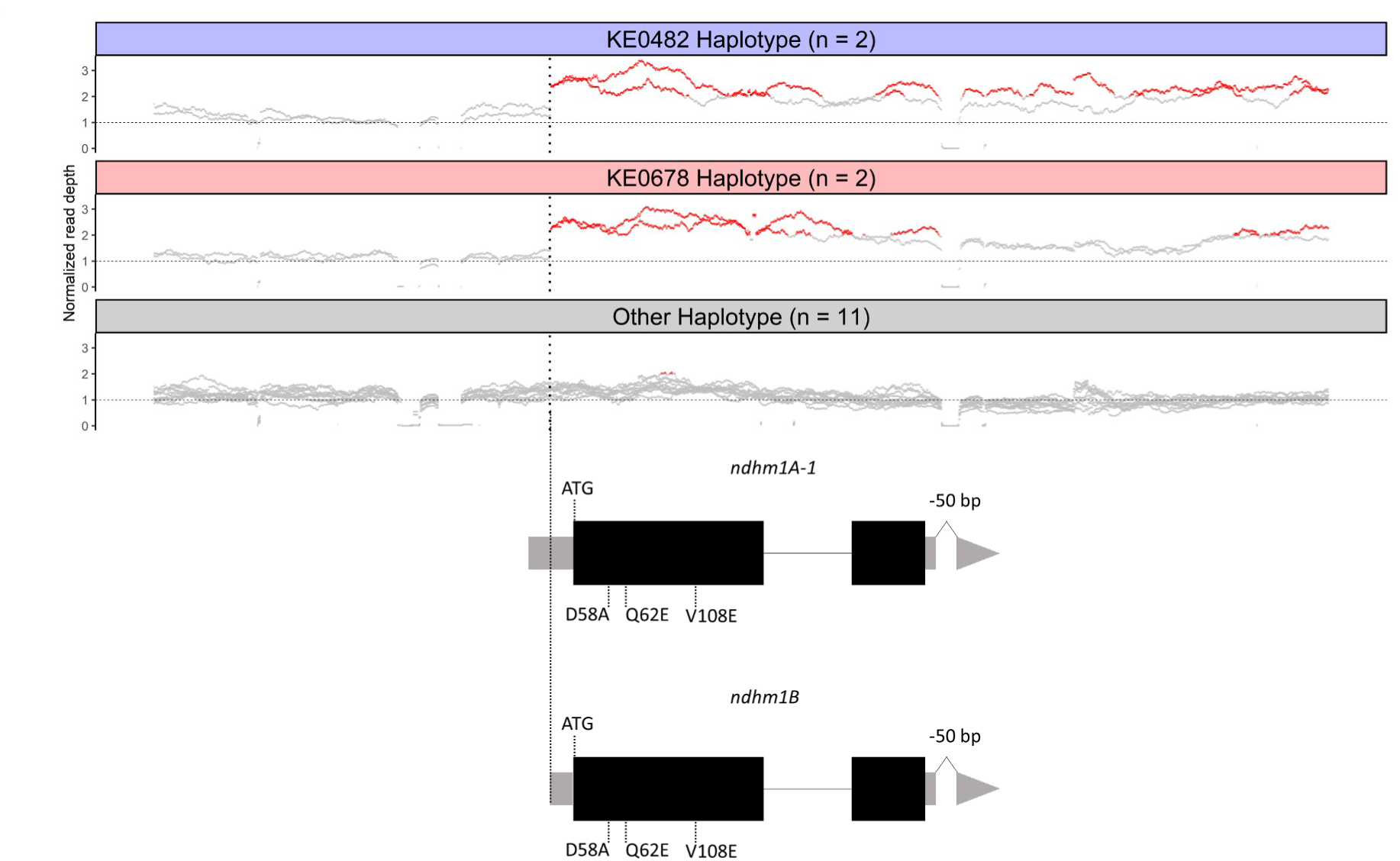
Association of lead haplotype alleles from GWAS of KE0482 and KE0678 with a duplication of *ndhm1*. Normalized read-depth of high-coverage WGS data aligned against B73v4 reference genome grouped by allele of the haplotype with lowest p-value from GWAS. Each line represents a Kemater line and each dot represents a single base in B73v4 reference. Hap1 and Hap2 are grouped together in the group “Other Haplotype”. Red: Normalized read-depth > 2.

### Effect of a transposon insertion in *ndhm1* on maize cold stress tolerance

EPH, Fv/Fm and chlorophyll content are connected to cold tolerance during early development through sensitivity of maize photosynthesis to cold temperatures (Burnett and Kromdijk, 2022; Lainé et al., 2023). Thus, we subjected Kemater lines with different *ndhm1* alleles to cold stress in controlled conditions. A subset of 15 Kemater lines was chosen based on previous results on allelic diversity of *ndhm1* and evaluated under optimal conditions and three days after recovery from severe cold stress (48 h, 6 °C / 2 °C [day/night]). Under optimal conditions, maximum and operating quantum efficiency of PSII (Fv/Fm, ΦPSII) as well as electron transfer rates (ETR) were significantly lower in lines with the *ndhm1A-1*(-/-)/*ndhm1B*(+/+) compared to lines with *ndhm1A-1(+*/+)/*ndhm1B*(-/-). In contrast, Fv/Fm, ΦPSII and ETR were increased in Kemater lines with the combination *ndhm1A-1*(+/+)/*ndhm1B*(+/+) compared to lines with ndhm1A-1(+/+)/ndhm1B(-/-) (Figure 7A, Figure S12). Lines with the hAT transposon insertion in *ndhm1A* were severely impaired in photosynthetic parameters three days after the cold stress, most prominent for ΦPSII and ETR, while the other lines recovered already (Figure 7A, Figure S12). After recovery, lines with *ndhm1A-2* showed necrosis on leaves, reflected in increased electrolyte leakage (Figure 7B, E).

**Figure 7:**
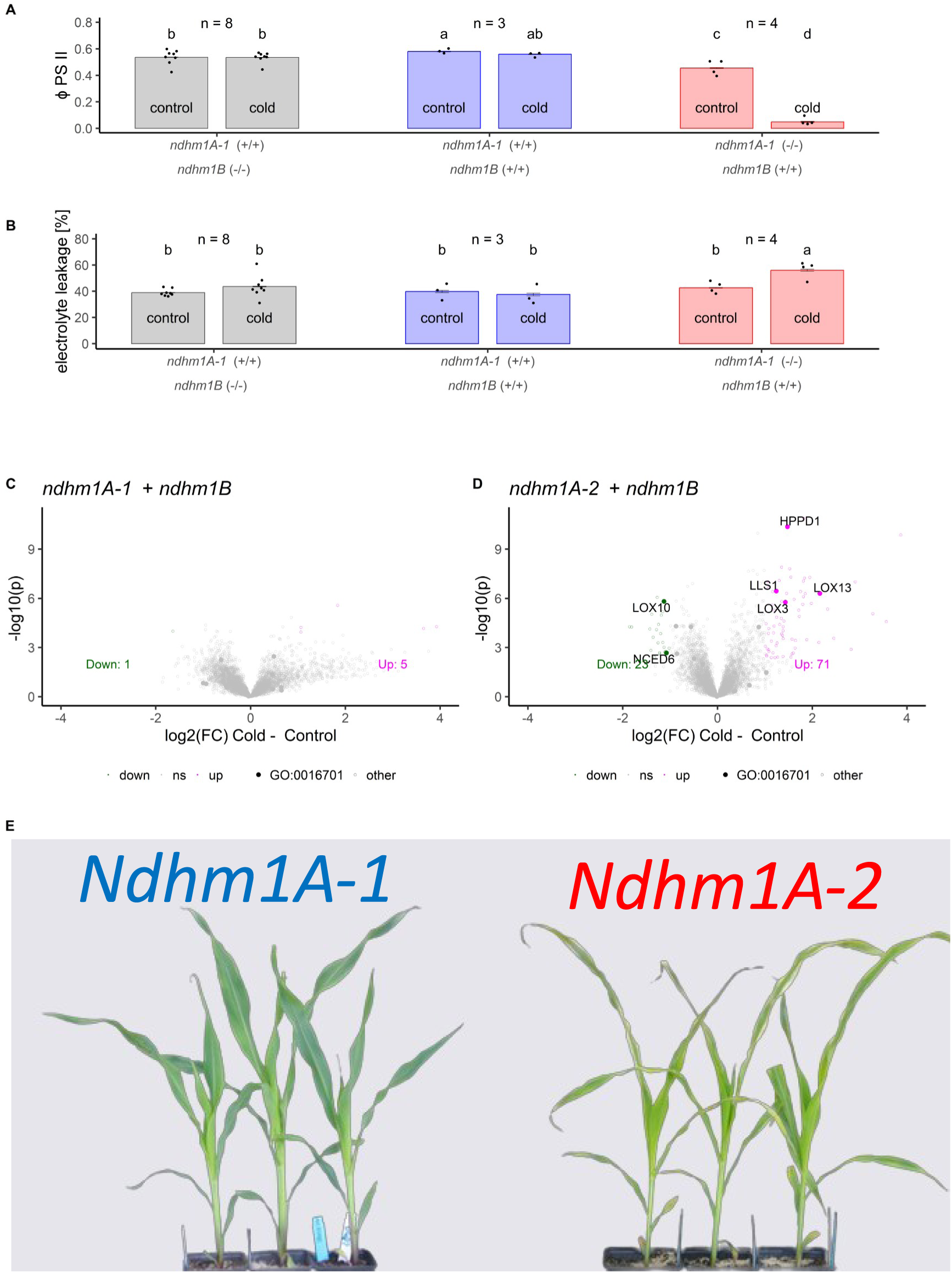
Effect of *ndhm1* allele after severe cold treatment on photosynthetic parameters and leaf proteome. A-B) Phenotypic differences of DH lines with different *ndhm1* alleles in optimal conditions and three days after recovery from severe cold stress. One point is the mean of 3 – 6 biological replicates of a Kemater DH line. Bars show means ± SE. Significant differences (lsd test) are indicated by letters. C, D) Difference of protein levels depending on a cold treatment (4°C / 2°C, 48 h) of genotypes with *ndhm1A-1* (C, KE0482, HIF1A, HIF3A) or *ndhm1A-2* (D, KE0678, HIF1B, HIF3B) alleles. One dot is one protein, filled dots are proteins with oxidoreductase activity (GO:0016701). Cut-off for differential expression: log2(FC) > 1; FDR <5%. Differentially expressed proteins with oxidoreductase are labeled in the plots. E) Lesions on leaves of Kemater lines KE0482 and KE0678 after cold treatment.

To investigate the effect of the transposon insertion in *ndhm1A* on cold stress tolerance we recorded leaf protein levels directly after a severe cold stress. In the leaf proteome of lines with *ndhm1A-1*, short term reactions to cold stress were almost absent (6 differentially accumulated proteins). In contrast, 94 proteins were differentially accumulated after cold treatment in lines with the hAT transposon insertion in *ndhm1A* (Figure 7C-D). Gene ontology enrichment revealed oxidoreductase activity (GO:0016701) as most significantly enriched molecular function in differentially accumulated proteins in lines with *ndhm1A-2*. Differentially expressed proteins with oxidoreductase activity include three lipoxygenases (LOX3, LOX10, LOX13) and two dioxygenases (HPPD1, NCED6) as well as one protein taking part in chlorophyll catabolism (LLS1).

## Discussion

Early maize planting offers both environmental and economic benefits to agriculture by reducing nitrogen leaching, minimizing herbicide use, avoiding summer drought during flowering, and enhancing yield potential through earlier soil coverage and extended growth periods. However, early maize planting is limited by low temperatures during early development (Kucharik, 2006; Parker et al., 2017; Lainé et al., 2023). Genetic variation for physiological responses to low temperatures has been described in maize, but the underlying genes are still unclear and only few candidate genes have been proposed (Reviewed in: Burnett and Kromdijk, 2022).

Landraces harbor genetic diversity crucial for improving traits with limited variation in breeding material (McCouch et al., 2013; Dwivedi et al., 2016; Mayer et al., 2020; Liang et al., 2021). The Austrian landrace Kemater was selected for this study due to its adaptation to temperate European climate and its variance for visible early growth traits such as EPH (Mayer et al., 2017; Hölker et al., 2019). Here, we demonstrate the concurrent genetic improvement of multiple quantitative traits crucial for enhancing early plant development through the action of the gene *ndhm1* (Fracheboud et al., 2004; Hund et al., 2004; Revilla et al., 2016; Enders et al., 2019). Variation of basic agronomic traits such as EPH as well as physiological parameters photosynthetic efficiency (Fv/Fm, ΦPSII), chlorophyll content and cold tolerance are directly linked to NDH-mediated cyclic electron transfer in our study. The multifaceted functionality of *ndhm1* underscores the pivotal role of NDH in maintaining metabolic processes critical for plant growth and development during early development.

Starting from a GWAS analysis we identified a hAT transposon insertion in *ndhm1* as the factor underlying a QTL with pleiotropic effects on the aforementioned traits due to its strong phenotypic consequences. NDH deficiency underlying the QTL was congruent with impaired NDH complex formation leading to specific chlorophyll induction curves, as well as reduced net CO_2_ assimilation rate, growth parameters and chlorophyll content in maize mutants of NDH subunits NDF6 and NDHU (Zhang et al., 2024). We observed unexpected Fv/Fm effects specific to *ndhm1* alleles in Kemater lines and a Mu transposon insertion mutant of NDHM in the genomic background of Flint line F7. This contrasts with earlier findings where NDH-deficient mutants did not impact Fv/Fm in C_4_ but exclusively in C_3_ species and only following cold or high-light stress (Endo et al., 1999; Wang et al., 2006; Zhang et al., 2024). The Fv/Fm effects might therefore be specific to defects in the assembly of subcomplex A, where NDHM is located, while NDF6 and NDHU are situated in subcomplex ED (electron donor) and subcomplex B, respectively (Shen et al., 2022; Zhang et al., 2024). Reduced Fv/Fm is indicative of photoinhibition of photosystem PSII and could explain enhanced PTOX2 protein levels (Calzadilla et al., 2024; Nie et al., 2024). However, in NADP-ME type photosynthesis, NDH is enriched in bundle sheath (BS) cells where PSII is mostly absent, which might suggest a distinct role of NDHM in NDH-mediated CET(Zhang et al., 2024).

Capitalizing on the native diversity of *ndhm1* in the landrace Kemater, we identified an allele which was associated with improved early development and photosynthetic traits. A haplotype allele of *ndhm1* was associated with a duplication of *ndhm1* and enhanced early development, photosynthetic efficiency and chlorophyll content. Noticeable candidates for the phenotypic effect of the improved allele are a hAT transposon insertion upstream of *ndhm1A* and a second copy of *ndhm1 (ndhm1B)*, however we cannot exclude the contribution of other, less striking polymorphisms without further studies. Our data indicate that *ndhm1B* is still transcribed to RNA, however deciphering the role of *ndhm1B* warrants more research and goes beyond the scope of this study. Thus, the mechanism underlying phenotypic variation associated with the improved haplotype remains somewhat elusive. Another important factor in function of the NDH complex is spatial expression in bundle sheath and mesophyll cells which is strictly regulated and dependent on the sub type of photosynthesis (Ishikawa et al., 2016). Evaluating the impact of allelic variation through cell-type-specific comparisons will be required to unravel the molecular mechanisms which are underlying the allelic effects of *ndhm1* we observe.

While genome editing continues to gain traction in crop improvement, the challenges in engineering alleles that truly enhance target traits with quantitative inheritance persist. As exemplified here, landraces offer an excellent platform for enhancing maize performance in breeding programs, particularly considering the regulatory complexities associated with biotechnological interventions (Liang et al., 2021; Lorenzo et al., 2023; Marone et al., 2023). Mapping of relevant genes influencing quantitative traits of interest is facilitated by the segregation of large effect QTL in landrace populations, followed by extensive analysis of allelic diversity to discover alleles that enhance target traits. The effect of the improved *ndhm1* allele we detected is relatively small compared to the defective allele but significant and impacting the same set of traits (EPH, Fv/Fm, ΦPSII, ETR and chlorophyll content). On molecular level, the improved allele exists in the US NAM panel. Further research will be needed to quantify its effect size in different genetic material. While NAM lines were chosen to be highly diverse we were able to evaluate distinct effects of three allelic classes of *ndhm1* in a homogeneous genomic background which allows us to connect allelic variation to meaningful phenotypes (Hufford et al., 2021).

Previous research linked genetic variation of CET to differences in salt and heat tolerance in C_3_ species soybean and rice (He et al., 2015; Essemine et al., 2017), but evidences are not conclusive and to our knowledge not available for C_4_ plants such as maize. Our research suggests potential for optimization of CET in maize breeding to increase EPH, Fv/Fm and chlorophyll content and provides an improved allele that could be directly used for breeding. We propose the utilization of maize landraces adapted to temperate climates as a valuable resource for accessing beneficial alleles aimed at enhancing early plant development and facilitating gene discovery efforts.

## Material and Methods

### Plant Material

Doubled-haploid (DH) lines extracted from the Austrian landrace “Kemater Landmais Gelb” (KE) and derived recombinants were used in this study. Detailed information about the generation, genotyping and phenotyping of the source material is described elsewhere (Hölker et al., 2019; Mayer et al., 2020; Mayer et al., 2022).

Two DH lines of the landrace Kemater (KE0482, KE0678) differed for multiple overlapping QTL on chromosome 2 for the traits early plant height (EPH), maximum potential quantum efficiency of PSII (Fv/Fm) and chlorophyll content (SPAD, Table S1) and were chosen for their similar genomic background (86,110 of 501,124 markers polymorphic, Figure S2). The two DH lines were crossed for generating a segregating F_2_ population, recombinant inbred lines (RILs) and heterogenous inbred families (HIFs) by consecutive selfing.

A subset of 27 out of 471 Kemater DH lines, carrying four different haplotypes at the QTL and selected to have an increased frequency of the KE0678- and KE0482-associated *ndhm1* haplotype was evaluated in the field (Validation set field) with a focus on early development. Another, partially overlapping subset of 16 Kemater DH lines carrying balanced frequencies of *ndhm1* alleles was evaluated in growth chambers (Cold growth chamber set) with a focus on recovery after cold stress.

We obtained Mu transposon insertion lines for the gene *ndhm1* from the BonnMu collection in the genomic background of the inbred line F7 (Marcon et al., 2020; www.bonnmu.uni-bonn.de; Win et al., 2024). F7 is an important founder of European Flint maize (Haberer et al., 2020) and was suitable because of its genomic similarity to the Kemater landrace.

### Development of recombinant inbred lines and heterogeneous inbred families

A schematic overview of material development is presented in Figure 1A. KASP markers positioned in the QTL region on chromosome 2 and polymorphic between the two parental lines (KE0482, KE0678) were synthesized using the probe sequences of the publicly available 600k Axiom™ Maize Genotyping Array (Thermo Scientific). RILs were genotyped and individual plants showing recombination between markers AX-91512997 (Chr2: 21380448, B73_AGPv4) and AX-90737994 (Chr2:29359872, B73_AGPv4) were self-pollinated. Resulting F_2:3_ RILs were phenotyped in a field experiment in three German locations in 2020.

RILs heterozygous for the target region were repeatedly self-pollinated and F_5_ plants genotyped using KASP markers (Table S5). Individual plants which carried a heterozygous fragment in the target region were self-pollinated to develop heterogeneous inbred families (HIFs). Heterogenous families are pairs of lines contrasting for a genomic region of interest derived from the same parental plant to enable phenotyping in near-isogenic backgrounds. To contrast the QTL effect in near isogenic backgrounds for fine-mapping resulting offspring was genotyped using KASP markers and the two homozygous classes (HIF1A, HIF1B, HIF3A, HIF3B) selected.

### Selection of Mu insertion mutants

Mu transposon insertion mutants in the genomic background of Flint line F7 were genotyped with *ndhm1* specific primers in combination with the mutant specific TIR6 primer (Figure S13, Table S6, "ndhm1 Exon1 fw and "ndhm1 5UTR rv" respectively; Settles et al., 2004). Amplification of respective gene fragments was verified by Sanger sequencing of the amplicons after separation by agarose gel electrophoresis and purification (NucleoSpin® Gel and PCR Clean-up kit, Macherey-Nagel GmbH & Co. KG, Düren, Germany). Individual plants were propagated to obtain homozygous mutants and wildtypes and were phenotyped for EPH, Fv/Fm and SPAD.

### Growth conditions

For phenotyping in growth chamber experiments kernels were imbibed in water for 5 min, transferred to filter paper and pre-germinated for 72 h in the dark at 28 °C. After germination, plant material was sampled from individual seedlings for DNA extraction and the respective seedlings were transferred to small pots filled with CL ED73 soil (Einheitserdewerke Patzer, Germany). Plants were grown until growth stage V4 to V6 with 16 h of light a day at 25 °C during the day and 20 °C during the night with 650 µmol m^-2^s^-1^ photosynthetically active radiation (PAR) at 75 % relative humidity (RH) for 6 biological replicates per DH line and treatment. In cold stress experiments plants were randomly assigned to a control group and a cold stress group which was treated with severe cold stress (6°C/2°C [day/night]) for 48 h, starting 18 days after sowing.

Field experiments were conducted in Einbeck (EIN, Germany, 51°49’05.9"N 9°52’00.3"E), Roggenstein (ROG, Germany, 48°10’47.5"N 11°19’12.9"E), Bernburg (BBG, Germany, 51°49’28.6"N 11°42’26.3"E), Oberer Lindenhof (OLI, Germany, 48°28’26.3"N 9°18’17.9"E) and Freising (FRS, Germany, 48°24’13.2"N 11°43’28.5"E) in the years 2020 and 2023. 178 F_2:3_ RILs were randomized in augmented block designs in EIN, ROG and BBG in 2020 with inbred checks UH007, F283, DK105 and EP44 replicated in each block. The subset of 27 DH lines used for molecular analysis of *ndhm1* allelic diversity in the field were randomized in complete blocks with two replications in locations ROG, FRS and OLI in 2023 using the same inbred checks. Plots consisted of 20 plants each in single rows of 3 m length with 0.75 m spacing between plots (9 plants/m^2^). Field trials were subjected to standard agricultural practices.

### Collection of agronomic traits

A detailed description of phenotypic data collection of the full set of Kemater DH lines is given by Hölker et al. (2019) and Mayer et al. (2022).

Early plant height was measured by stretching all leaves of a plant to measure the maximum length between soil and the tip of the leaves. The measurements of three to five individual plants per plot were averaged to obtain the plot level measurement. Days to silking and days to pollen shedding were calculated as number of days from sowing until half of a plot had silks of at least 1 cm length or until half of a plot shed pollen, respectively.

### Determination of photosynthetic traits

Fv/Fm was measured in dark adapted plants before light was turned on in the growth chambers using a LI-600 porometer (LI-COR Inc., Lincoln, NE, USA). In field experiments, plants were measured after midnight to ensure that plants were dark adapted for at least one hour. The last fully developed leaf was clipped in the middle, omitting the midvein, flow rate set to ‘high’ (150 µmol*s^-1^), match frequency of 10, flash set to ‘dark adapted’ and ‘Rectangular’ with an intensity of 6000 µmol m^-2^s^-1^, a flash length of 800 ms, and the fluorescence constants ‘Leaf absorptance’ and ‘Fraction Abs PSII’ to 0.8 and 0.5 respectively at a modulation rate of 5 Hz. ΦPSII and ETR were measured on the same day at noon after adapting the plants for several hours to the light conditions in the growth chamber using the LI-600 with adapted settings. The flash intensity increased to 10000 µmol m^-2^s^-1^ the setting ‘Dark Adapted’ was disabled and the actinic modulation rate was set to 600 Hz.

Chlorophyll content was measured using a SPAD-502 (Konica Minolta K.K, Chiyoda, Tokyo, Japan), clipping the last fully developed leaf of a plant. Data from three plants per plot were averaged to obtain the plot-level measurement.

For NDH-mediated CET fluorescence induction curves were recorded on dark adapted plants in the morning before the light was turned on in the growth chambers with the LI-6800 (LI-COR Inc., Lincoln, NE, USA). The last fully developed leaf was clipped with the measuring light (ML) set to 50 Hz and a weak actinic light (AL) applied (50 µM m^2^s^-1^) for 5 minutes per plant until the fluorescence signal remained stable. After reaching a steady state level the ML was turned off and the fluorescence signal recorded for two more minutes.

CO_2_ response curves of net photosynthetic assimilation rate (A/Ci) were recorded as described by Blankenagel et al. (2022) in field grown plants before flowering at ambient CO_2_ concentrations of 0, 50,100, 200, 300, 400, 600, 800, 1000, 1200 ppm.

### Transcript level measurements

RNA was extracted from leaves using a CTAB protocol (Logemann et al., 1987), followed by DNase digestion and first-strand cDNA synthesis (Maxima H Minus Kit, random hexamer primers, Thermo Scientific K1652). RT-qPCRs were performed in technical triplicates with primers binding to the 3’ UTR of *ndhm1* (Table S6). For normalization, RT-qPCR was performed for the house-keeping gene MEP (membrane protein PB1A10.07c, *Zm00001d018359*). In field experiments leaf tips of three plants per plot and two replicates in location FRS were harvested at noon in growth stage V6 and pooled in one sample per genotype.

### Proteomics experiment

For proteome measurements HIF1A, HIF1B, HIF3A, HIF3B, KE0482 and KE0678 were grown in a growth chamber and cold treated as described above. The last fully developed leaf was sampled directly after the cold stress in frozen nitrogen for three biological replicates per genotype – treatment combination. Total proteome was extracted and measured following an established protocol (Brajkovic et al., 2023) with an additional step of 11-plex TMT labeling of peptides after protein digestion for quantification. For normalization of batch effects, a reference was prepared by mixing equal peptide amounts of all samples which was measured twice in each of the TMT batches. Proteins were identified and quantified by MaxQuant (Tyanova et al., 2016) using B73_AGPv4 as reference followed by rescoring the identified proteins using Prosit (Gessulat et al., 2019). To avoid imputation of missing values in quantitative analysis, only proteins which were identified in all 36 samples (6 Genotypes x 3 Replicates x 2 Treatments; N = 5667 Proteins) were considered. Log2 transformed raw intensities of identified proteins were median normalized to remove differences in summed intensities per sample and subsequently the two reference channels per TMT batch were used to calculate a correction factor for each identified majority protein group per TMT batch. Gene ontology term enrichment was performed by agriGO v2.0 (Tian et al., 2017) comparing differentially expressed proteins to all 5667 proteins quantified in the proteomics experiment as custom background.

### Sample preparation for mass spectrometry

The dried and cleaned peptides were reconstituted in 20 µl of 200 mM EPPS (pH 8.5) buffer. TMT-11plex reagent (Thermo Scientific) was reconstituted in water-free ACN to a working concentration of 20 µg/µl. 5 µl of this TMT reagent solution was transferred to the peptides. The reaction was incubated on the thermoshaker (20°C, 400 rpm) for one hour and quenched by 0.25 % hydroxylamine afterward. The TMT channels were then pooled together and acidified with FA to a final concentration of 1 %. The reaction wells were washed with 20 µl washing solution (10 % ACN, 10 % FA) and added to the TMT pool. The TMT pools were dried down in the speed-vac and stored at -20°C. The 1 mg TMT-pooled peptides were cleaned by solid-phase extraction on 50 mg C18 Sep-PAK cartridges. The TMT peptides were washed with 0.1 % FA. TMT peptide elution was achieved by 0.1 % FA in 60 % ACN. TMT peptides were dried down in the speed-vac and stored at -20°C. The sample was reconstituted in 25 mM ammonium bicarbonate and was fractionated on a Waters XBridge BEH C18, 4.6×250 mm over a 44 min gradient from 7 to 45 % ACN in the presence of 2.5 mM ammonium bicarbonate. After gradient, a 6 min ramp to 80 % ACN followed to wash the column. Fractions were collected from minutes 7 to 55. Each fraction consisted of 30 sec at a flow rate of 1 ml/min. The 96 fractions were pooled back into 48 fractions in an n-with-(n+48) fashion. Samples were acidified with FA to a final concentration of 0.1 %. The fractionated TMT peptides were dried down in the speed-vac and stored at -20°C until MS measurement.

### Mass Spectrometry

Full proteome TMT-labeled peptides were measured with a Fusion Lumos Tribrid mass spectrometer (Thermo Scientific) that was coupled to a Dionex Ultimate. The sample was directly injected onto the Acclaim PepMap 100 C18 column (2 µm particle size, 1 mm ID × 150 mm). Separation was performed on a 27 min segmented gradient with a flow rate of 50 µl/min starting from 4% to 27% B (23min) and 27% to 32% B (2min). The system was finally washed with 100 μl 90 % B and re-equilibrated at 1 % B. Solvent A consisted of 0.1 %v FA and 3 %v DMSO in water. Solvent B consisted of 0.1 %v FA and 3 %v DMSO in Acetonitrile.

The MS was operated in a fast, data-dependent MS3 -mode. The spray voltage was set to 3.5 kV supported by sheath gas (32 units) and aux gas (5 units) with a vaporizer temperature of 125 °C. Every 1.2 s, a full-scan (MS1) was recorded from 360 to 1600 m/z with a resolution of 60k in the Orbitrap in profile mode. The MS1 AGC target was set to 4e5. Based on the full scans, precursors were targeted for MSMS scans if the charge was between 2 and 6, the isotope envelop was peptidic (MIPS), and the intensity exceeded 1e4. The MS2 quadrupole isolation window was set to 0.6 Th. The TMT peptides were HCD fragmented with an NCE of 34 %. The MS2 spectra were acquired in the ion trap in rapid mode. The MS2 AGC target was set to 3e4 charges, and the maxIT was set to 40 ms. The maxIT or AGC target could be dynamically exceeded when the previous scan took longer than the calculated injection time (inject beyond mode). TMT reporter ions were measured in a consecutive MS3 scan based on the previous MSMS scan. Thus, a new batch of precursor ions was isolated with an MS3 quadrupole isolation window of 1.2. The isolated precursor was then HCD-fragmented identically to the previous MS2 scan. Additionally, Isobaric tag loss exclusion properties were set to TMT reagent. The selected fragment ions were then HCD fragmented with an NCE of 55 %. The MS3 spectrum was acquired with 50k resolution from 100 to 1000 Th in the Orbitrap in centroid mode. The MS3 AGC target was set to 2e5 charges, and the maxIT was set to 86 ms.

### Detection of different *ndhm1* alleles

To identify different *ndhm1* alleles, we designed primers for the full-length variant (*ndhm1A*) and for the variant with shortened 5’UTR (*ndhm1B,* Figure 2C, Figure S13). The primers “ndhm1 Exon 2 fw” and “ndhm1 5UTR rv” were used for full length *ndhm1* transcript detection. For *ndhm1B*, “Genotyping copy unspecific fw”, “Genotyping copy unspecific rv” were used. Genotyping for the transposon insertion in *ndhm1A-2* was done by discriminating amplicon lengths of transcript or genomic sequences by gel electrophoresis. All primers sequences are listed in Table S6. PCR reactions were performed using Q5 High-Fidelity DNA Polymerase (New England Biolabs Inc., Ipswich, MA, USA) according to the manufacturer’s manual at an annealing temperature of 68 °C for 15 s followed by elongation at 72 °C for 30 s and PCR products separated by gel electrophoresis.

### Sanger sequencing

Sanger sequencing was performed using Mix2seq kits at Eurofins Genomics Germany GmbH (Ebersberg, Germany).

### Whole-genome sequencing

Kemater DH lines KE0095, KE0109, KE0482, and KE0678 representing three different *ndhm1* haplotype alleles were sequenced using PacBio HiFi long reads by CNRGV, INRA Occitanie Toulouse, France (cnrgv.toulouse.inra.fr). Circular consensus sequences were de-novo assembled using the hifiasm assembler with default parameters (Cheng et al., 2021). The contigs were ordered using a genetic map derived from EP1xPH207 as reference in ALLMAPS (Tang et al., 2015; Haberer et al., 2020).

For fine-mapping, HIF1A, HIF1B, HIF3A, HIF3B, KE0482 and KE0678 were sequenced with high coverage (50 X) using Illumina short-reads. Paired-end short reads were de-duplicated (hts_SuperDeduper v1.0), trimmed (Trimmomatic v0.39), error corrected using Kmer distributions (Lighter v1.1.2) and mapped against the KE0678 PacBio assembly using bwa-mem (Li and Durbin, 2009). SNPs were called from resulting bam files of HIF1A, HIF1B, HIF3A, HIF3B, KE0482 and KE0678 by freebayes (Garisson and Marth, 2012). Settings for variant calling were: –ploidy 2, –min-mapping-quality 1, –min-base-quality 3, –min-alternate-count 5 –min-repeat-entropy 1, –no-partial-observations –no-population-priors –genotype-qualities. Variants were subsequently filtered by bcftools using following parameters: FORMAT/DP > 10; INFO/DP < 100, %QUAL > 30, SAF > 0 & SAR > 0, RPL >1 & RPR > 1. Heterozygous SNPs in the DH lines KE0482 and KE0678 were removed as no heterozygosity is expected in DH lines. Finally, SNPs differentiating KE0678, HIF1B and HIF3B from KE0482, HIF1A and HIF3A were selected and used to define the fine-mapped genomic segment in KE0678.

For read-depth analysis 15 Kemater DH lines (column “WGS”) were sequenced with high coverage (50 X) using Illumina short-reads. Paired-end short reads were de-duplicated (hts_SuperDeduper v1.0), trimmed (Trimmomatic v0.39), error corrected using Kmer distributions (Lighter v1.1.2) and mapped against the B73_AGPv4 reference, carrying a single-copy of *ndhm1*, using bwa-mem (Li and Durbin, 2009; Jiao et al., 2017). Read-depth of 15 Kemater DH lines per position of B73_AGPv4 reference was extracted using the command ‘samtools depth’ from BAM files. The coverage per position was normalized by dividing by the mean read depth over the whole B73_AGPv4 reference for each genotype.

### Comparative genomic analysis

Genomic positions of KASP markers in the target regions of KE0482 and KE0678 were obtained by mapping their probe sequences against KE0482 and KE0678 PacBio HiFi assemblies genomes using bwa-mem (Li and Durbin, 2009). Genomic positions of B73_AGPv4 gene features (Jiao et al., 2017) in KE0482 and KE0678 were identified by BLAST.

Pairwise sequence alignment of the target region between KE0482 and KE0678 was conducted using nucmer with the setting -c 156, -g 1000 and subsequently filtered using delta-filter with settings -i 99 and -l 1000 (Kurtz et al., 2004).

A MOASeq binding site partially overlapping with the 5’UTR of *ndhm1* in B73_AGPv4 was identified and downloaded from MaizeGDB (https://jbrowse.maizegdb.org/?loc=chr2%3A23629472..23634058&tracks=gene_models_official%2Cmoa_seq_coverage_peaks&highlight=) using the track ‘MOA-seq coverage peaks’ and searched by BLAST against the Kemater PacBio assemblies (Savadel et al., 2021).

The exon structure of *ndhm1* alleles was determined by PCR amplification of cDNA fragments using exon spanning primers followed by Sanger sequencing and re-aligning the obtained sequences against the assembly of KE0678 using BLAST.

### Statistical Analysis

All statistical analyses were done in R (R Core Team, 2013). For field experiments with Kemater DH lines adjusted means were estimated for each trait using the R package ‘ASReml-R’ (v4.1.0). The statistical model for calculating adjusted means was

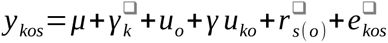

where µ is the overall mean; γ_k_ is the fixed effect of DH line k; u_o_ is the random effect of environment o; γu_ko_ is the random interaction effect for genotype k and environment o; r_s(o)_ is the random effect of the replicate s (nested in environments) and e_kos_ is the residual error.

Strong phenotypic segregation between the F_2:3_ RILs of leaf color and early plant height and required scoring of plots as “KE0482-like”, “KE0678-like” or “segregating” based on leaf color and early plant height. Linkage of genome-wide KASP markers with the “KE0482-like” and “KE0678-like” phenotype was calculated by chi-square goodness of fit tests for each KASP marker individually followed by correction for multiple testing. To determine the mode of gene action of *ndhm1* in F_2:3_ RILs the closest marker to *ndhm1* (AX-90736551) was used to calculate the mean of homozygous RILs for KE0482 and KE0678 allele or being heterozygous, respectively. The mean of both homozygous classes was compared to the mean of heterozygous RILs by a two-sided Student’s t-test.

For significance testing in fine-mapping and mutant analysis, two-sided Student’s t-tests were conducted between KE0482 and KE0678, HIF1A and HIF1B, HIF3A and HIF3B and the Mu mutant and its wildtype, respectively, for EPH, Fv/Fm, SPAD and RT-qPCR data. Differences in CO_2_ assimilation rates were assessed by fitting a mixed linear model for each ambient CO_2_ concentration using the genotypic score of the *ndhm1* allele as fixed effect and the genomic background (HIF1, HIF3) as random effect. Significance of allele effects was assessed by Wald tests (Kenward and Roger, 1997). For the phenotypic comparisons of multiple *ndhm1-*associated haplotypes in field and growth chamber experiments least significant differences (lsd) were applied.

For proteome analysis two-sided Student’s t tests were calculated for the difference of protein amounts between *ndhm1* alleles for each protein separately. To avoid false positive associations, proteins were considered significantly differentially expressed at a false discovery rate of 5% (Benjamini and Hochberg, 1995) and a log2 fold change bigger than one.

## Acknowledgements

We thank Brigitte Neuhauser, Iris Prücklmaier, Sylwia Schepella, Stefan Schwertfirm and Margot Siebler for technical assistance. We thank the Plant Technology Center (Technical University of Munich, Germany) for providing infrastructure and technical support during greenhouse, growth chamber and field experiments. We would like to acknowledge the support of William Marande and Caroline Callot from CNRGV, INRAE (http://cnrgv.toulouse.inrae.fr/) for providing assistance in PacBio long read sequencing and GENTYANE platform of Clermont-Ferrand INRAE Center (http://gentyane.clermont.inra.fr/) for providing access to PacBio sequencer.

## Author contributions

C.-C.S., V.A., M.O., P.W. and S.U. designed the research and developed ideas; S.U., V.A. and B.O. designed and performed phenotyping experiments and analyzed the data; S.R. and S.U. analyzed whole genome sequencing data; C.M. and F.H developed Mu transposon mutants; S.B. acquired and analyzed proteomics data. K.E. and S.U. conducted candidate gene analysis; S.U., M.M., M.O., T.P., D.S. and C.U. developed the plant material; S.U, V.A., P.W. and C.-C.S. wrote the manuscript; all authors read and approved the final manuscript; C.-C.S. agrees to serve as the author responsible for contact and to ensure communication.

---

**Table S1:** QTL detected for EPH and photosynthetic traits in landrace Kemater. Regions were defined using 10-SNP haplotype windows as described in Mayer et al. 2020 and adjusted means of the respective phenotype across up to 11 environments from Hölker et al. 2019. In the column “Effect size (sd)” the minimal and maximal effect sizes within an environment of the most significantly associated haplotype expressed as standard deviations of the respective trait is indicated. Genomic positions are given in B73_AGPv4 coordinates and N genes indicates the number of gene models within these regions based on B73_AGPv4 coordinates. Start- and end haplotype indicate the genomic positions of the most significantly associated haplotype within a QTL.

| Trait | Chr | Start QTL | End QTL | Start Haplotype | End Haplotype | Effect size (sd) | N genes | Size (kb) |
| --- | --- | --- | --- | --- | --- | --- | --- | --- |
| EPH (V6) | 2 | 22942795 | 25023002 | 23199379 | 23288438 | 0.14-0.24 | 60 | 2080 |
| Fv/Fm (V4) | 2 | 21191092 | 25675167 | 23328380 | 23333363 | 0.18-0.35 | 121 | 4484 |
| Fv/Fm (V6) | 2 | 23083888 | 23333363 | 23328380 | 23333363 | 0.18-0.32 | 8 | 249 |
| SPAD (V3) | 2 | 22974990 | 23333363 | 23328380 | 23333363 | 0.21-0.33 | 9 | 358 |

**Table S2:** Candidate genes in 314 kb fine-mapped QTL region on chromosome 2.

| **Chr** | **Start** | **End** | **B73v4 ID** | **Functional Annotation** |
| --- | --- | --- | --- | --- |
| 2 | 23362233 | 23364607 | Zm00001d002811 | C2C2-GATA transcription factor |
| 2 | 23397758 | 23400810 | Zm00001d002812 | rmr2 - required to maintain repression 2 |
| 2 | 23401605 | 23402088 | Zm00001d002813 |  |
| 2 | 23462897 | 23483867 | Zm00001d002814 |  |
| 2 | 23483486 | 23484612 | Zm00001d002815 | NDHM |
| 2 | 23485039 | 23486175 | Zm00001d002816 |  |

**Table S3.** Frequencies of most associated 10-SNP haplotype from GWAS found in Kemater DH population (N = 471). Hap3 and Hap4 represent the alleles of the parents of the bi-parental population used for fine-mapping the QTL. In the columns N (PacBio) and N (WGS 50X) the genomic datasets used in this study for the respective haplotype is indicated.

| Haplotype | N | Allele Frequency | N (PacBio) | N (WGS 50 x) |
| --- | --- | --- | --- | --- |
| Hap1 | 206 | 43.7 % |  | 6 |
| Hap2 | 133 | 28.2 % | 2 | 5 |
| Hap3 (KE0678) | 102 | 21.7 % | 1 | 2 |
| Hap4 (KE0482) | 30 | 6.4 % | 1 | 2 |

**Table S4.** Genotyping of *ndhm1A-1* and *ndhm1B* in a set of 27 Kemater lines for in depth molecular characterization of allelic diversity.

| **Haplotype** | **N Lines** | ***ndhm1A-1*** | ***ndhm1B*** |
| --- | --- | --- | --- |
| Hap1 | 3 | 3 | 0 |
| Hap2 | 3 | 3 | 1 |
| Hap3 (KE0678) | 15 | 3 | 14 |
| Hap4 (KE0482) | 6 | 6 | 6 |
| **Total** | **27** | **15** | **21** |

**Table S5:** Genomic positions of KASP markers used to genotype RILs and HIFs on chromosome 2 of different genome assemblies. Positions were identified by mapping the probes of the publicly available Axiom™ Maize 600k Genotyping Array against the respective genome assemblies using bwa-mem. Genomic positions of B73 refer to B73_AGPv4 coordinates.

| Marker | Chromosome | Position KE0482 | Position (KE0678) | Position (B73) |
| --- | --- | --- | --- | --- |
| AX-91512997 | Chr2 | 21786495 | 22402152 | 21380448 |
| AX-90574469 | Chr2 | 22397136 | 23228550 | 22168417 |
| AX-91513237 | Chr2 | 22831689 | 23639123 | 22578878 |
| AX-90736227 | Chr2 | 23344555 | 24137631 | 23114217 |
| AX-90736268 | Chr2 | 23565056 | 24244586 | 23330240 |
| AX-90736476 | Chr2 | 24458646 | 25197545 | 24006284 |
| AX-90736551 | Chr2 | 24865533 | 25522080 | 24392802 |
| AX-91513731 | Chr2 | 25285469 | 26152105 | 25023871 |
| AX-90737128 | Chr2 | 26669764 | 27271153 | 26327957 |
| AX-90737994 | Chr2 | 30769334 | 30380132 | 29359872 |

**Table S6:**
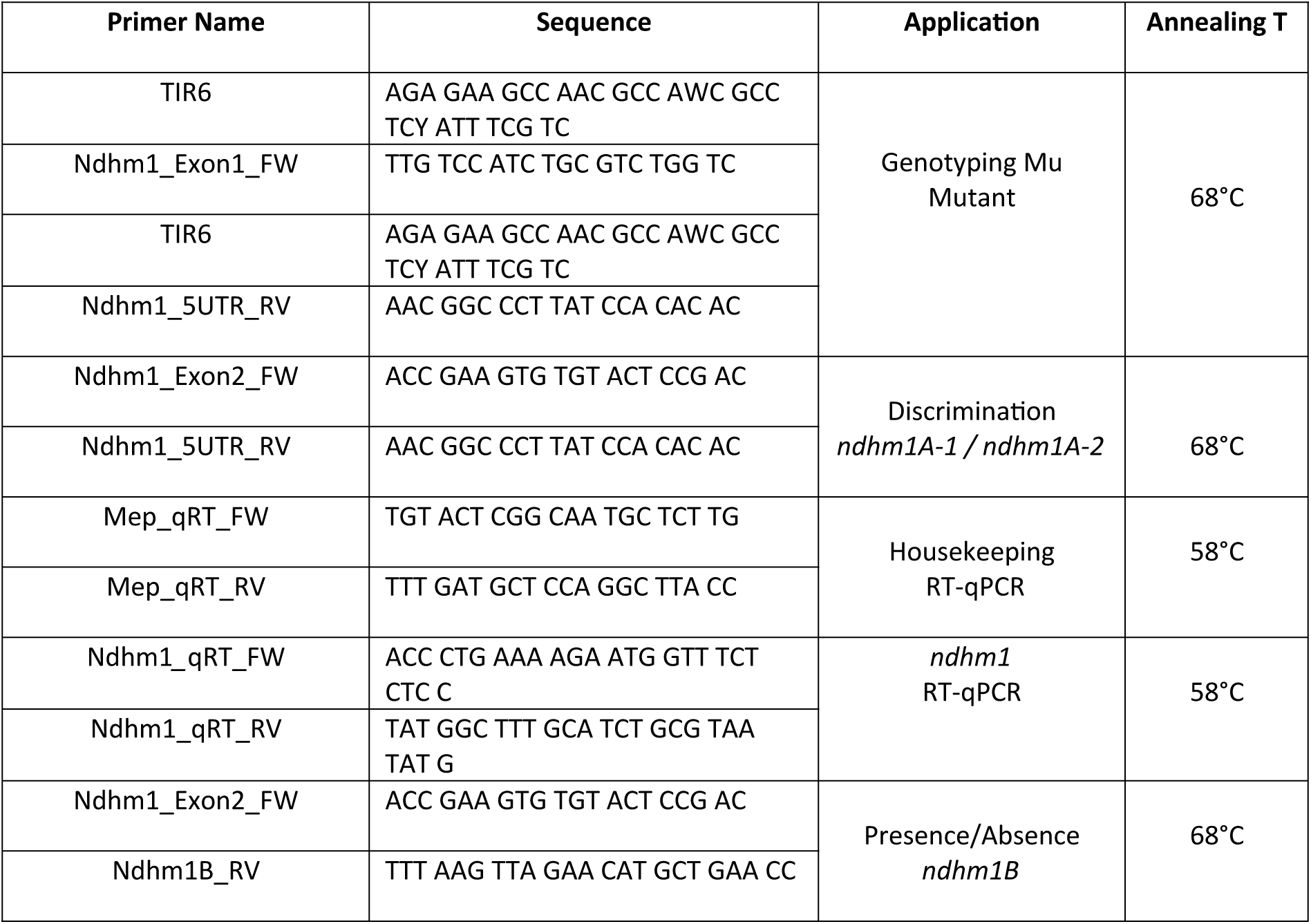
Primers used for RT-qPCR analysis and genotyping different *ndhm1* alleles. Elongation time was 30 sec for the PCRs and 60 sec for RT-qPCRs,

**Figure S1:**
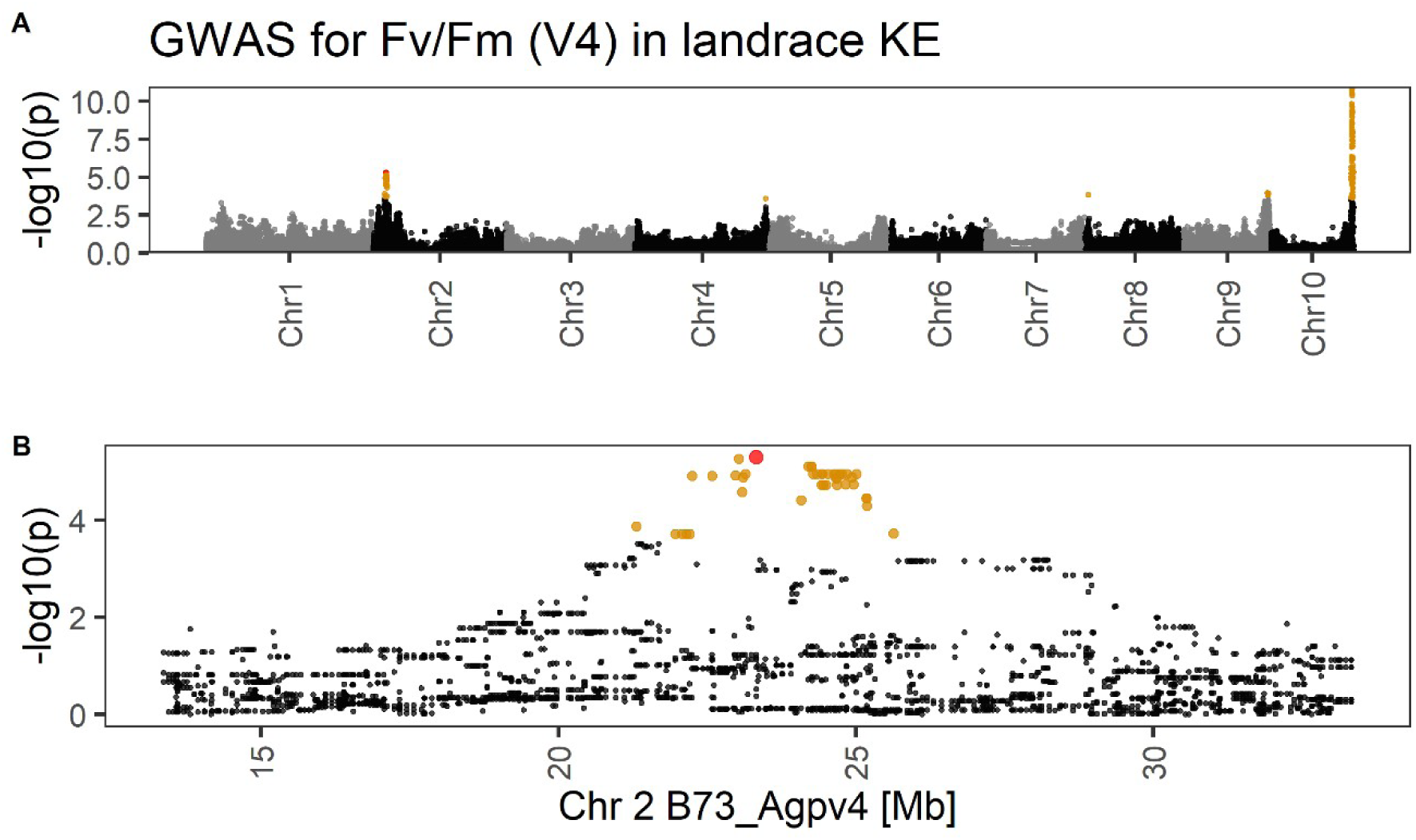
GWAS in Kemater DH lines. A) Manhattan plot for a haplotype-based GWAS for Fv/Fm following Mayer et al. 2020 within landrace KE (V4). Yellow: Haplotypes with FDR < 15 %. B) Zoom in the QTL on chromosome 2. Red: haplotype with the strongest association with Fv/Fm (lead haplotype).

**Figure S2:**
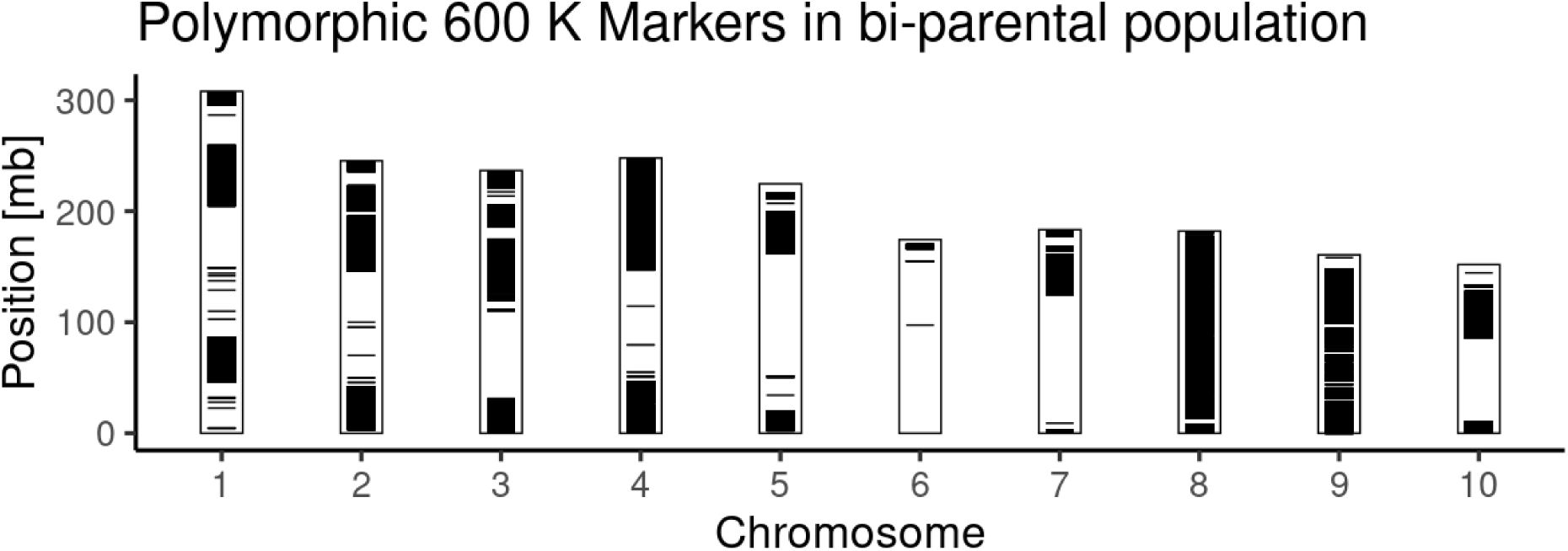
Polymorphic 600k Axiom™ Maize Genotyping Array SNPs between KE0482 and KE0678. The 600k dataset and filtering is described in detail in Hölker et al. 2019. In total 501,124 markers remained after quality control from which 86,110 (17 %) are polymorphic (black) and 415,014 are monomorphic (white) between DH lines KE0482 and KE0678.

**Figure S3:**
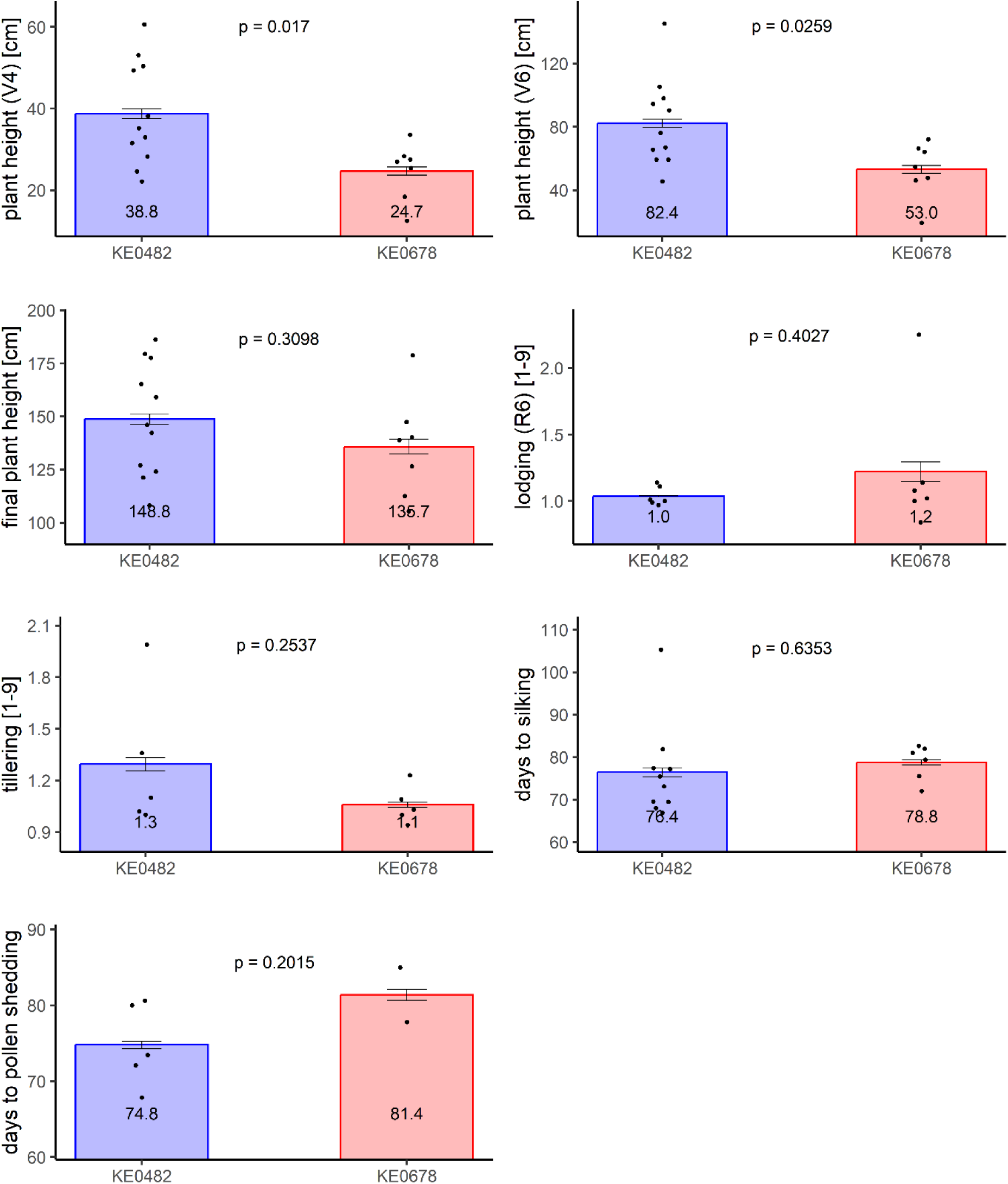
Phenotypic differences between DH lines KE0482 and KE0678 in field experiments in up to 11 combinations of locations and years (Hölker et al. 2019). Bars show means ± SE and dots adjusted means within one location-year combination. P-values are plotted above bars and were derived from a two-sided Student’s t test comparing the adjusted means between KE0482 and KE0678. The mean of within location-year adjusted means is plotted in bars.

**Figure S4:**
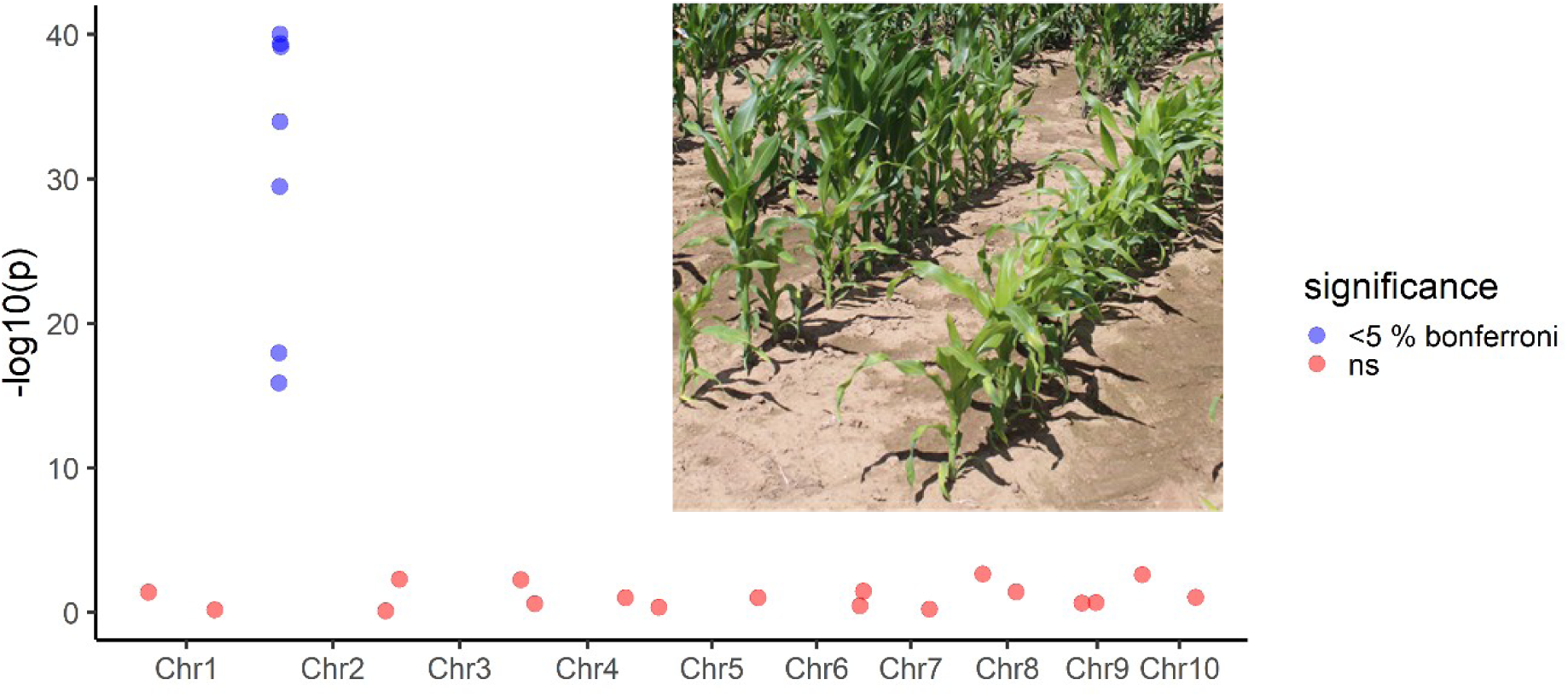
Chi-Squared test of independence of KE0678-like phenotype with genome wide KASP markers in 211 F_2:3_ RILs. F_2:3_ RILs were grown in 3 locations in 2020 and plots scored for KE0482-like or KE0678–like phenotype based on the low early vigor and plant height and leaf color of KE0678 (Figure S3, Picture below). For each marker a Chi-squared test of independence was performed. Bonferroni correction was applied to correct for multiple testing and markers with an adjusted p-value < 0.05 were considered to be dependent on the marker genotype.

**Figure S5:**
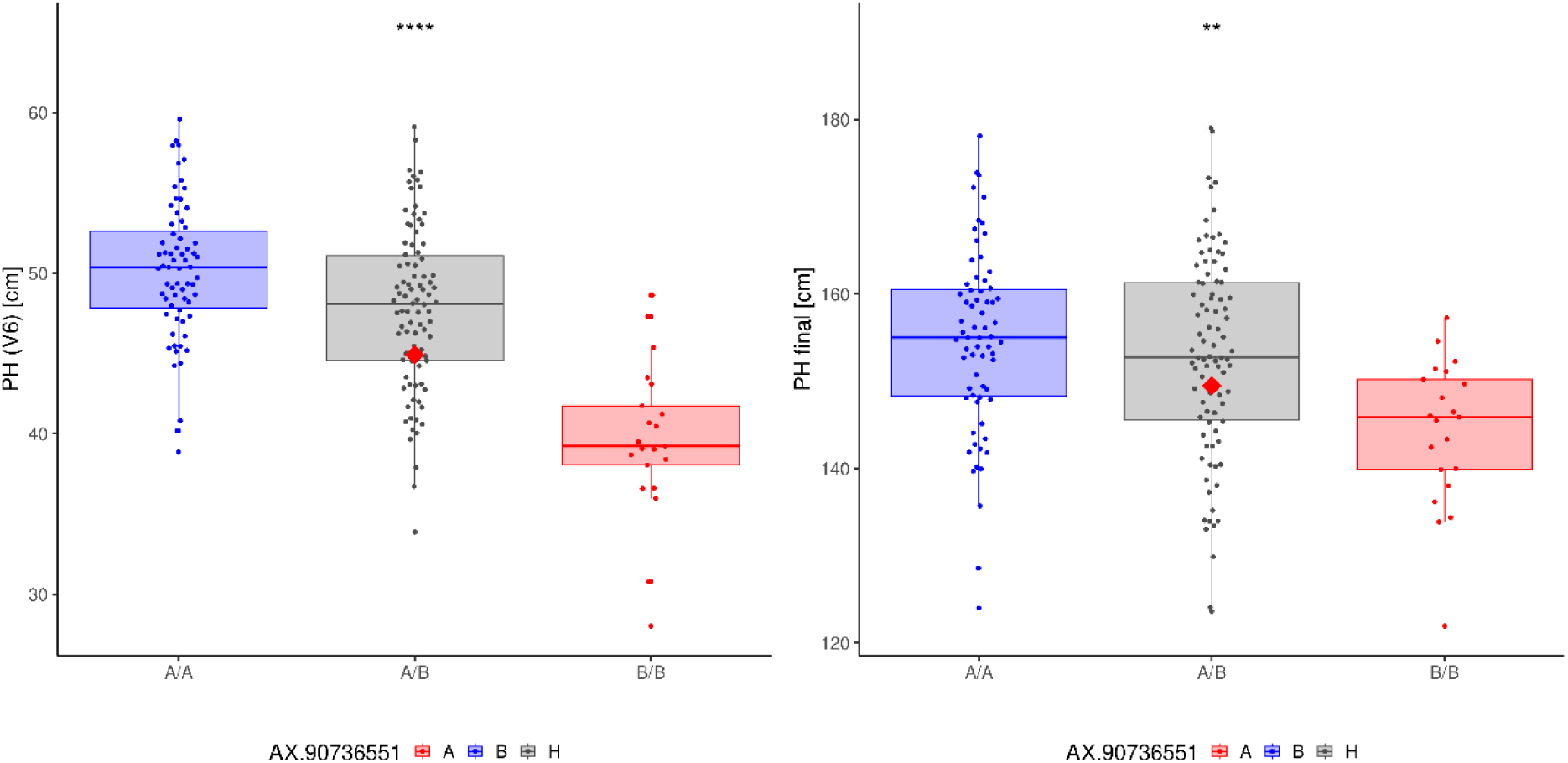
Test for deviation from midparent value in field experiment in 2020. Each data point represents an adjusted mean of one F_2:3_ recombinant inbred line (RIL) across three locations. Phenotypic data is grouped by the genotypic marker closest to *ndhm1* (AX-90736551). The mean of genotypic groups is indicated by a horizontal line in the boxplots. Allele A refers to the KE0482 allele, Allele B to the KE0678 allele and A/B to the heterozygous state. Red diamond: midparent value. Significant differences (t-test) between the midparent value and the mean of the A/B group are marked with stars. \*\*\**P* < 0.001, \*\**P* < 0.01.

**Figure S6:**
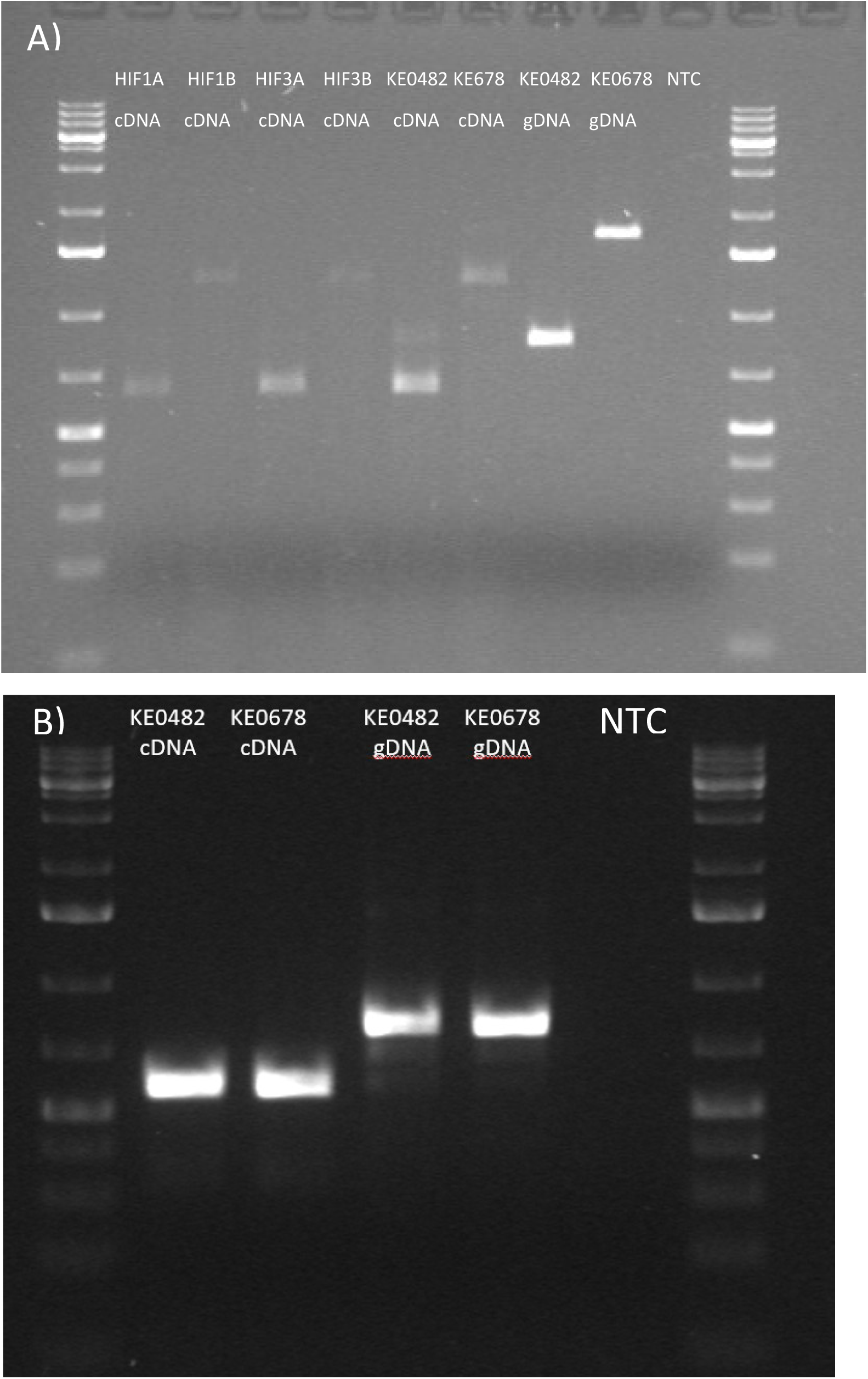
Transcription of *ndhm1* alleles in bi-parental population. Sequences of primers are listed in Table S6. As positive controls fragments from genomic DNAs were amplified from KE0482 and KE0678. A) Primers specific for *ndhm1A*. HIF: Heterogeneous inbred families. B) Primers specific for *ndhm1B*. NTC: No template control; gDNA: genomic DNA; cDNA: complementary DNA

**Figure S7:**
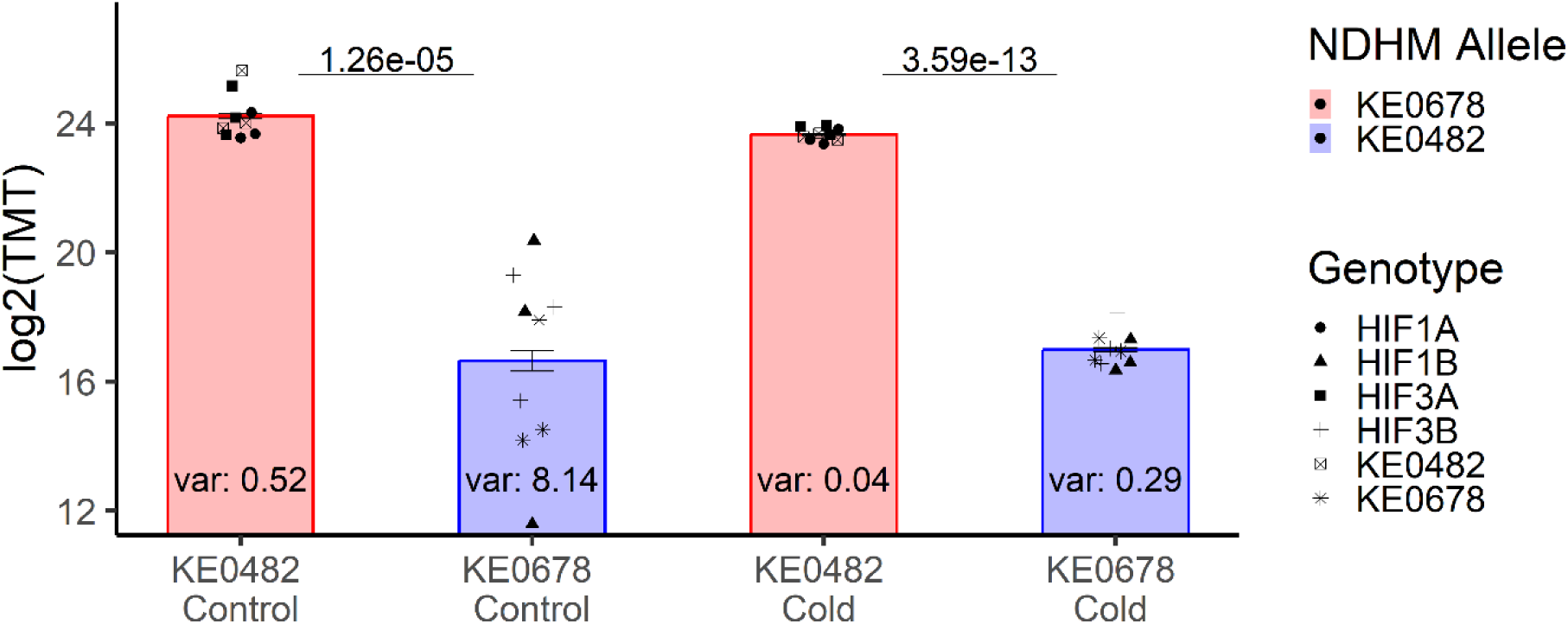
Differences in NDHM protein levels between KE0482, HIF1A, HIF3A and KE0678, HIF1B and HIF3B (N=3 per genotype). Bars show means ± SE (n = 9 plants) and dots NDHM protein amount from one replication of a genotype. Significance was assessed by a two-factor ANOVA with the NDHM allele and genomic background as factors independently for the both treatments. Variance within a group is indicated in the bars.

**Figure S8:**
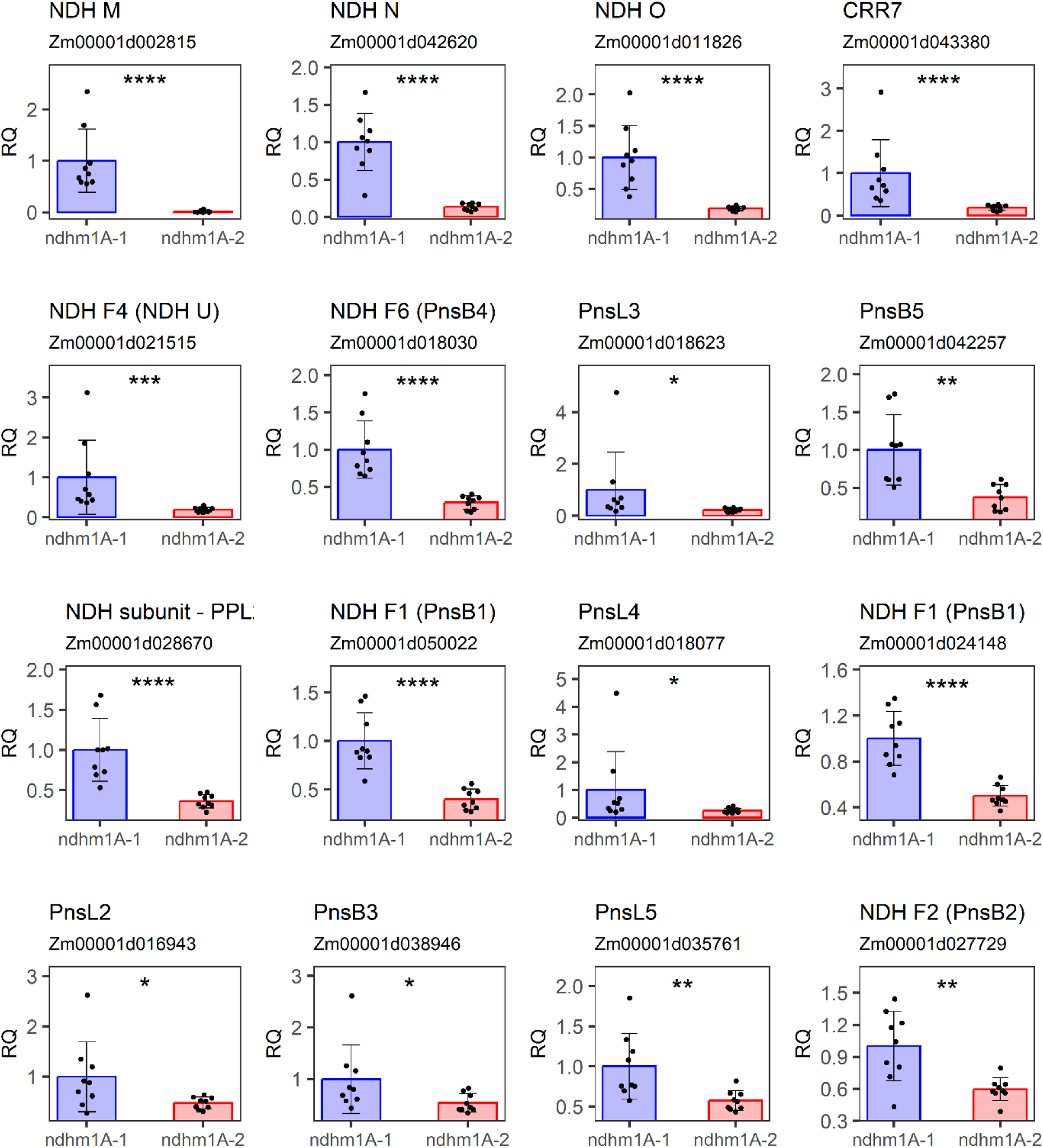
Impact of hAT insertion in *ndhm1A-2* on relative protein levels of NDH components. Bars show means ± SE (n = 9 plants) and dots relative NDHM protein amount from one replication of a genotype. Significant differences (t-test) are marked with stars. **** P<0.0001, *** P<0.001, ** P<0.01, * P<0.05.

**Figure S9:**
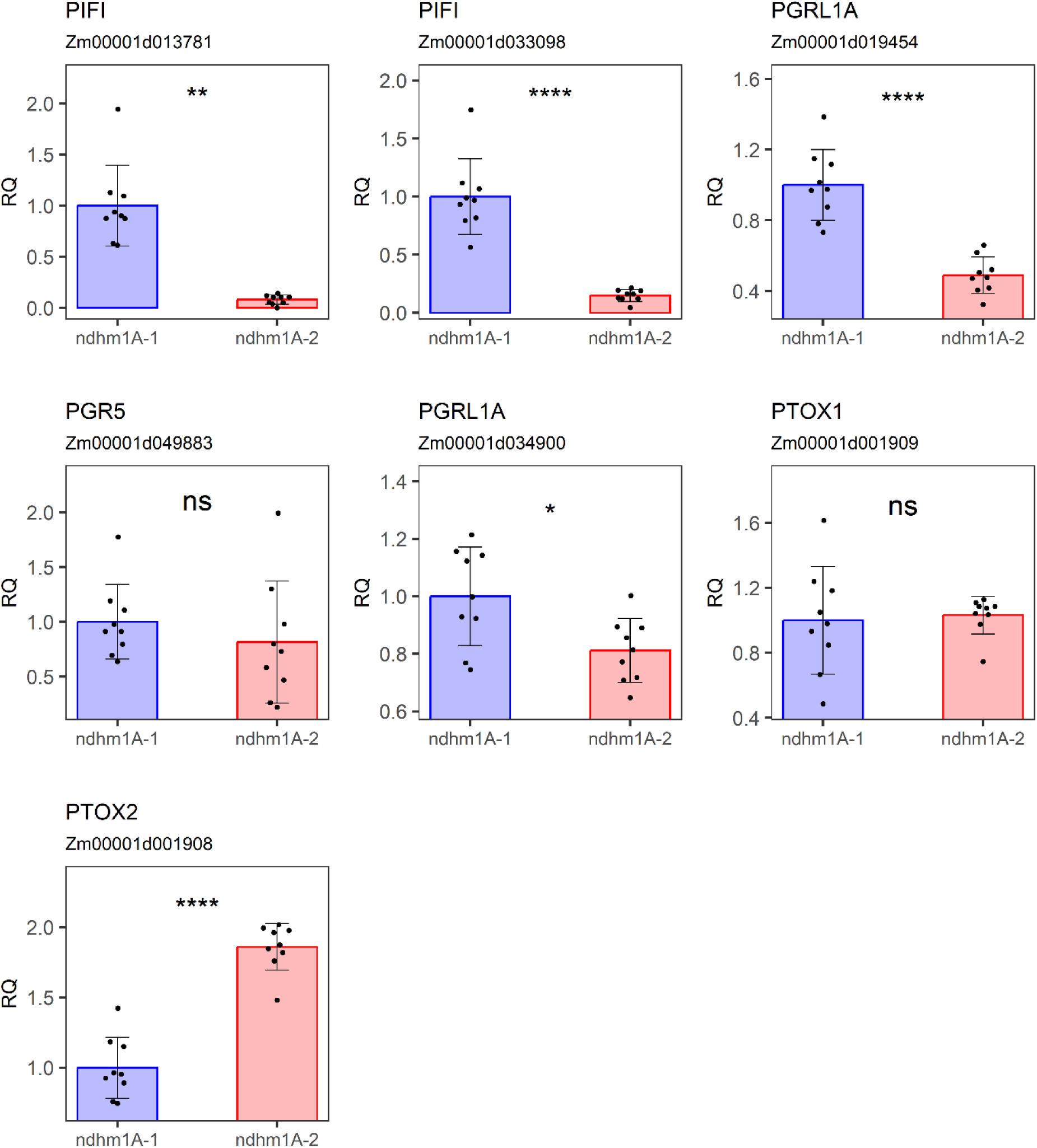
Impact of hAT insertion in *ndhm1A-2* on relative protein levels of non-NDH CET components. Bars show means ± SE (n = 9 plants) and dots relative NDHM protein amount from one replication of a genotype. Significant differences (t-test) are marked with stars. **** P<0.0001, *** P<0.001, ** P<0.01, * P<0.05, ns = not significant

**Figure S10:**
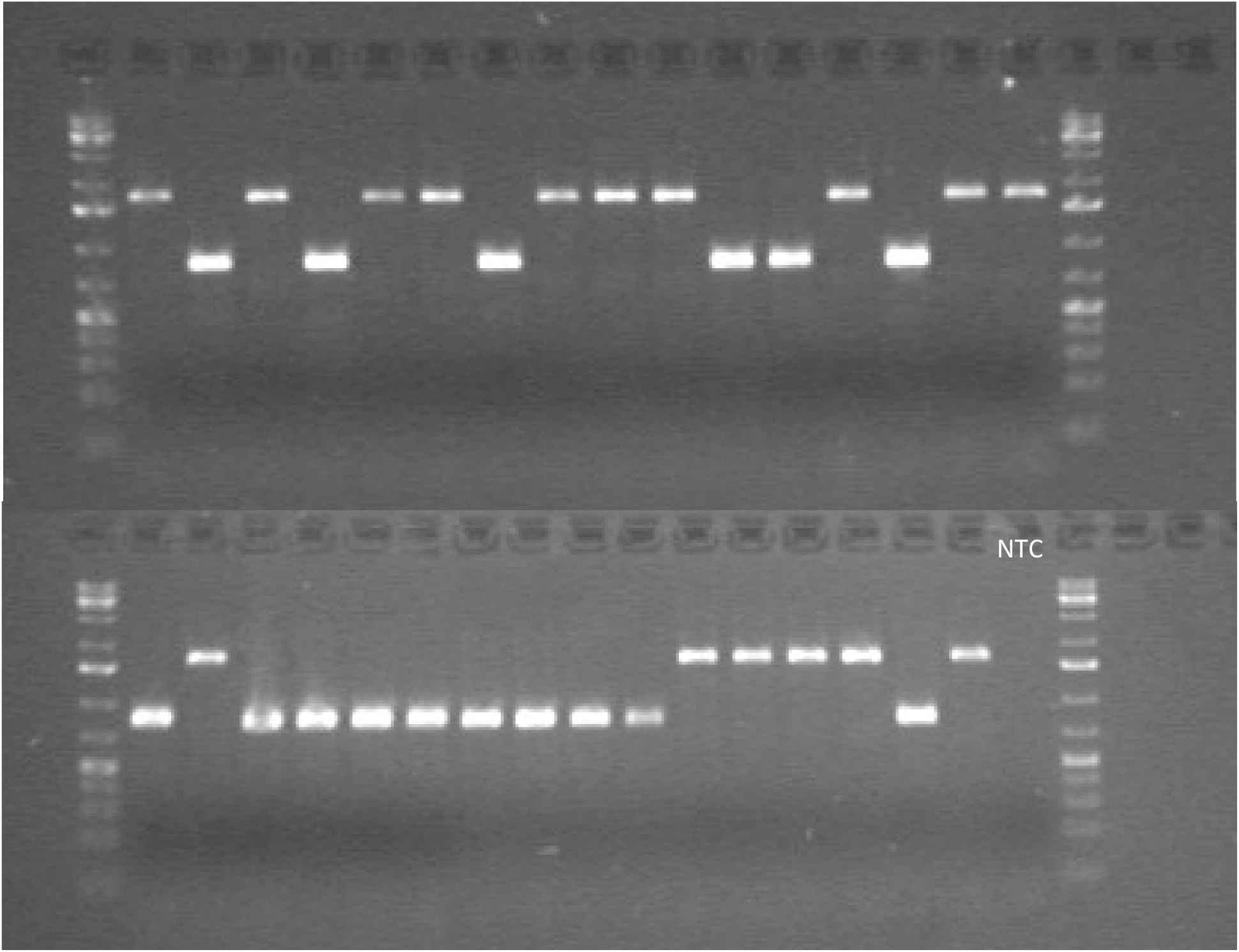
Genotyping of transposon insertion in *ndhm1A-2* in 27 Kemater lines. Lines with the transposon insertion display a longer DNA fragment in the gel. The first 27 samples are genomic DNAs of Kemater DH lines: KE0007, KE0011, KE0045, KE0054, KE0067, KE0079, KE0096, KE0103, KE0134, KE0153, KE0190, KE0194, KE0202, KE0204, KE0241, KE0277, KE0288, KE0294, KE0407, KE0426, KE0462, KE0482, KE0557, KE0590, KE0627, KE0633, KE0678. Samples 28-30 come from genotyping of recombinant HIF1B and samples 31 and 32 are internal reference genomic DNAs from KE0482 and KE0678, respectively. NTC: No template control.

**Figure S11:**
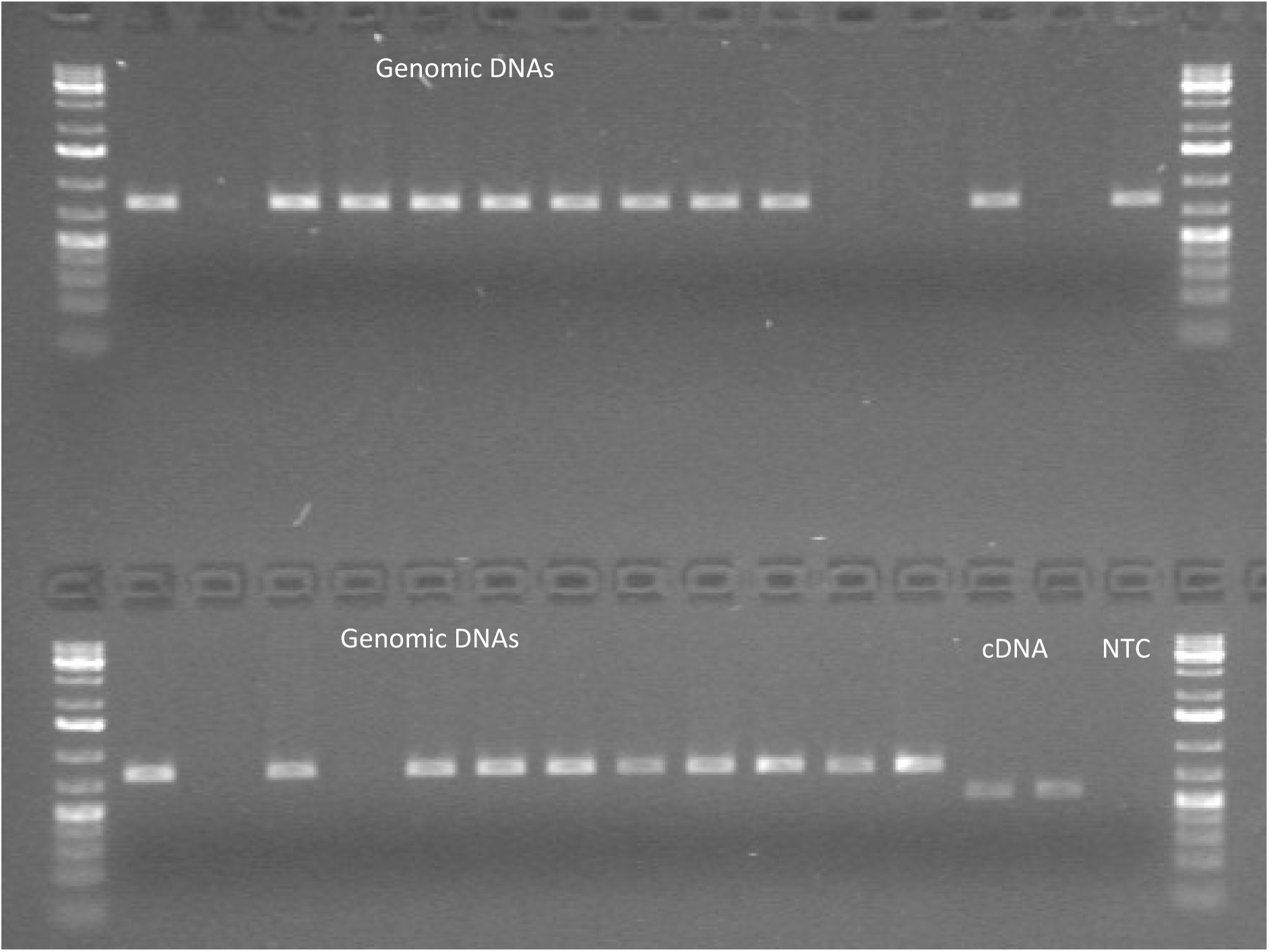
Genotyping of presence or absence of *ndhm1B* in 27 Kemater lines. The first 27 samples are genomic DNAs of Kemater DH lines: KE0007, KE0011, KE0045, KE0054, KE0067, KE0079, KE0096, KE0103, KE0134, KE0153, KE0190, KE0194, KE0202, KE0204, KE0241, KE0277, KE0288, KE0294, KE0407, KE0426, KE0462, KE0482, KE0557, KE0590, KE0627, KE0633, KE0678. The same primers were used to amplify cDNA from KE0482 and KE0678 with a shorter fragment length as the intron is spliced out. NTC: No template control; cDNA: complementary DNA

**Figure S12:**
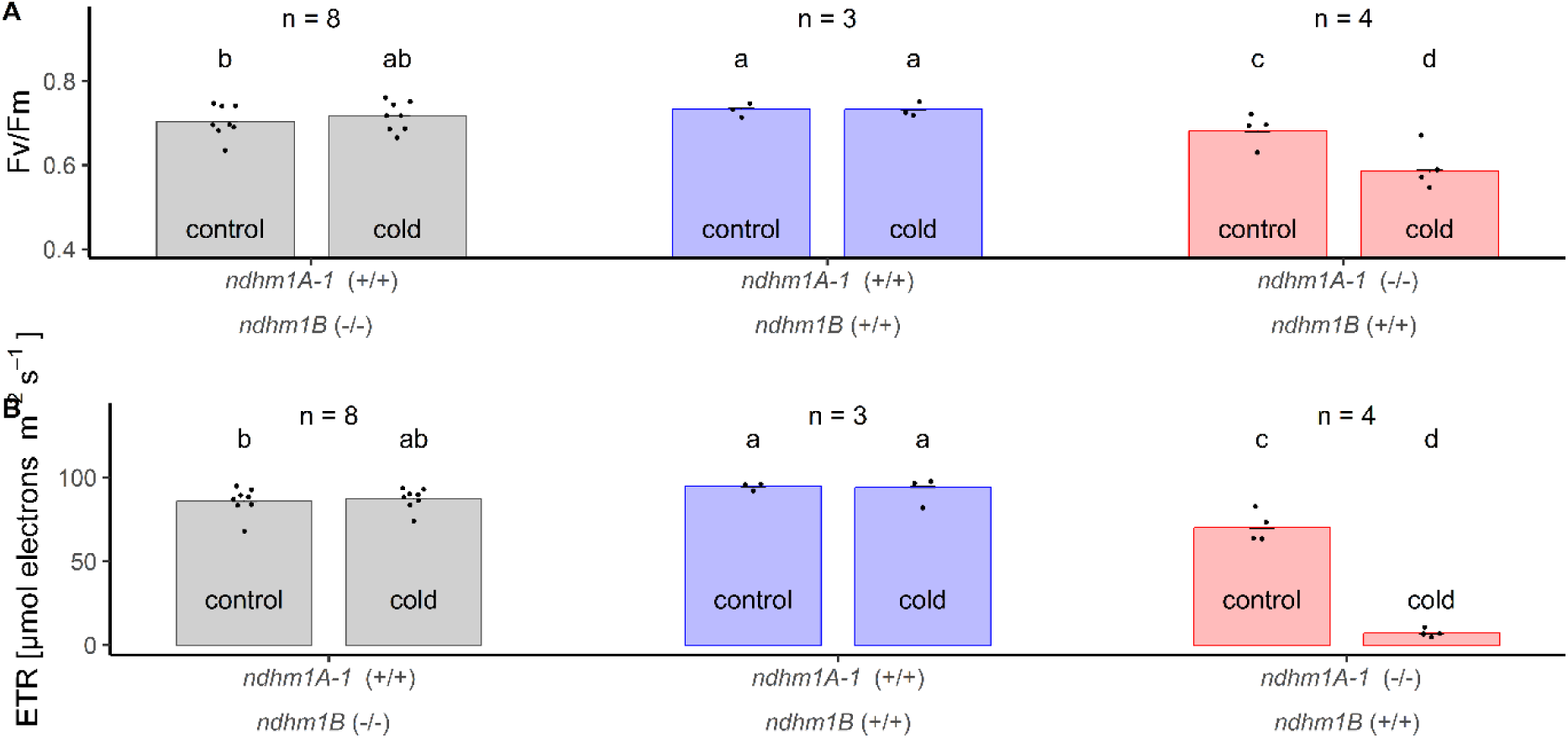
Effect of *ndhm1* allele after severe cold treatment on photosynthetic parameters. A-B) Phenotypic differences of DH lines with different *ndhm1* alleles in optimal conditions and three days after recovery from severe cold stress. A) Fv/Fm: Maximum potential quantum yield of PSII. B) ETR: Electron transfer rate. One point is the mean of 3 – 6 biological replicates of a Kemater DH line. Bars show means ± SE. Significant differences (lsd test) are indicated by letters.

